# Beyond the VSG Layer: Exploring the Role of Intrinsic Disorder in the Invariant Surface Glycoproteins of African Trypanosomes

**DOI:** 10.1101/2023.12.18.572128

**Authors:** Hagen Sülzen, Alexander N. Volkov, Rob Geens, Farnaz Zahedifard, Benoit Stijlemans, Martin Zoltner, Stefan Magez, Yann G.-J. Sterckx, Sebastian Zoll

**Affiliations:** Institute of Organic Chemistry and Biochemistry of the Czech Academy of Sciences, Flemingovo namesti 542/2, 16000, Prague, Czech Republic; Faculty of Science, Charles University, Albertov 6, 12800, Prague 2, Czech Republic; VIB-VUB Center for Structural Biology, Flemish Institute of Biotechnology (VIB), Pleinlaan 2, 1050 Brussels, Belgium; Jean Jeener NMR Centre, Vrije Universiteit Brussel (VUB), Pleinlaan 2, 1050 Brussels, Belgium; Laboratory of Medical Biochemistry (LMB) and the Infla-Med Center of Excellence, Department of Pharmaceutical Sciences, Universiteit of Antwerp, B-2610 Wilrijk, Belgium; Department of Parasitology, Faculty of Science, Charles University in Prague, Biocev, 25250 Vestec, Czech Republic; Brussels Center for Immunology (BCIM), Department of Bioengineering Sciences, Vrije Universiteit Brussel, B-1050 Brussels, Belgium; Myeloid Cell Immunology Laboratory, VIB Center for Inflammation Research, B-1050 Brussels, Belgium; Department of Biochemistry and Microbiology, Ghent University, B-9000 Gent, Belgium; Laboratory for Biomedical Research, Department of Molecular Biotechnology, Environment Technology and Food Technology, Ghent University Global Campus, Incheon 406-840, South Korea

## Abstract

In the bloodstream of mammalian hosts, African trypanosomes face the challenge of protecting their invariant surface receptors from immune detection. This crucial role is fulfilled by a dense, glycosylated protein layer composed of variant surface glycoproteins (VSGs), which undergo antigenic variation and provide a physical barrier that shields the underlying invariant surface glycoproteins (ISGs). The protective shield’s limited permeability comes at the cost of restricted access to the extracellular host environment, raising questions regarding the specific function of the ISG repertoire. In this study, we employ an integrative structural biology approach to show that intrinsically disordered membrane-proximal regions are a common feature of members of the ISG superfamily, conferring the ability to switch between compact and elongated conformers. While the folded, membrane-distal ectodomain is buried within the VSG layer for compact conformers, their elongated counterparts would enable the extension beyond it. This dynamic behavior enables ISGs to maintain a low immunogenic footprint while still allowing them to engage with the host environment when necessary. Our findings add further evidence to a dynamic molecular organization of trypanosome surface antigens wherein intrinsic disorder underpins the characteristics of a highly flexible ISG proteome to circumvent the constraints imposed by the VSG coat.

## INTRODUCTION

As strictly extracellular parasites, African trypanosomes rely on sophisticated defense, communication, and sensing mechanisms to evade host immune responses and initiate developmental transformations. Well-documented examples of the parasite’s bloodstream form (BSF, predominant form in the mammalian host) include: i) inhibition of innate immune responses through trypanosome receptor-like adenylate cyclases [1–3]; ii) complement and trypanolytic factor binding proteins [4–7]; iii) antigenic variation of variant surface glycoproteins (VSGs) to counter adaptive immune responses [8,9]; iv) inter-parasite communication through quorum sensing [10,11]; and v) the release of extracellular vesicles to manipulate the host environment, mediate parasite-parasite communication and facilitate specific forms of motility [12,13]. Among these, the most studied (and most notorious) mechanism remains the structure-function relationship of the parasite’s VSG coat. The classical model depicts the VSG coat as a dense, protective barrier that shields underlying invariant proteins from recognition by the host’s immune system. While small molecules may penetrate this shield, large immune reactive molecules at the size of an antibody have been shown to be excluded [14–17]. This viewpoint was further maintained by an extended model of the VSG coat organization, according to which VSG dimers can adopt two different conformations, generating a flexible topology while still maintaining the shielding function [18]. This, for the first time, linked the plasticity of a trypanosome surface protein to a specific function. However, as trypanosomes reside freely in the bloodstream, they must engage with numerous host ligands in the extracellular environment through dedicated surface receptors. The current VSG umbrella model and the assumed exclusion limit of the VSG coat raise questions as to how such surface receptors would be able to perform their biological function.

Invariant surface glycoproteins (ISGs) are a unique super-family of surface antigens that are exclusively expressed by the BSFs of African trypanosomes. Two distinct features set ISGs apart from other characterized BSF surface antigens. First, while most invariant receptors are confined to the flagellar pocket to minimize immune exposure [19,20], ISGs are embedded within the VSG coat and thus distributed over the whole cell surface [20–22]. Second, instead of being attached to the plasma membrane via GPI-anchors, ISGs are type-I transmembrane proteins with a short cytosolic domain, which is subject to ubiquitylation, therefore mediating rapid recycling [21,23]. Both features suggest a potentially important role in the immunobiology of BSFs, such capturing/scavenging nutrients or protecting the parasite from host immune factors. While several multi-gene ISG families have been identified, only ISG65 and ISG75 have been studied extensively due to their high abundance, indicating functional significance. Notably, ISG65 has recently been identified as a complement receptor [6,7,24], featuring an adaptation of the canonical three-helical bundle fold with unique characteristics, whereas experimental evidence for structure and biological function of ISG75 remain elusive. Owing to their relative abundance and invariant nature, ISG65 and ISG75 have been considered as potential vaccine candidates [22], as they are immunogenic and elicit an immune response detectable through the presence of antibodies in patient blood. Indeed, studies have shown that ISG75, ISG65, and ISG64 rank among the most abundant surface antigens and can be selectively enriched through immuno-affinity chromatography using *T. b. gambiense* infection IgG [25]. However, immunization with ISGs conferred no or only partial protection against repeated infections [26]. The accessibility of ISGs for the interaction with extracellular host ligands (including antibodies) becomes enigmatic should they strictly conform to the prevailing VSG shielding model [18,27]. Consequently, ISG exposure is likely to occur either through shedding or increased permissiveness of the VSG coat. Recent studies involving ISG65 from BSFs and metacyclic invariant surface proteins (MISP) from metacyclic forms have favored the latter hypothesis [7,28]. For both proteins, intrinsically disordered C-terminal regions (IDRs) were found, which could enable them to operate within as well as beyond the VSG coat. Protrusion beyond the coat’s boundaries would allow for the acquisition of large-sized host ligands at the expense of increased exposure to the host’s immune system. Despite this functional importance, the precise molecular mechanisms by which IDRs facilitate these phenomena remain unexplored.

In this study, we investigate the conformational flexibility of prototypical ISGs of different sizes from the human-infective parasite *T. b. gambiense* (*Tbg*ISG43, *Tbg*ISG64, and *Tbg*ISG75) and describe our findings in the context of the VSG umbrella model. Using an integrative structural modelling approach, combining AlphaFold2-based structure prediction, hydrogen-deuterium exchange mass spectrometry (HDX-MS), and small-angle X-ray scattering (SAXS)-driven conformational sampling, we demonstrate that, similarly to the previously described *Tbg*ISG65 [7], *Tbg*ISG43, *Tbg*ISG64, and *Tbg*ISG75 all possess disordered C-terminal linkers that are extremely variable in length. We show that these linkers have the capacity to adopt distinct conformational states, enabling the proteins to either reside within the VSG coat or protrude from it, thereby enabling them to interact with large extracellular ligands. Our results suggest that large-scale conformational changes mediated by intrinsically disordered membrane-proximal linkers are a functional feature of trypanosome receptors of the ISG super-family that may have evolved as an adaptation to their presence within the VSG coat.

## MATERIAL AND METHODS

### Protein production and purification

*T. b. gambiense ISGs*. ISG75 was cloned, recombinantly produced and purified as previously described in [29]. DNA fragments (Genewiz) encoding amino acids (aa) 22–349 of *Tbg*972.5.200 (from here on referred to as *Tbg*ISG43) and aa 24-365 from *Tbg*ISG64 (*Tbg*972.5.6550) were codon-optimized for bacterial expression and cloned into the pET15b plasmid using the Gibson assembly method (New England Biolabs).

*Tbg*ISG43 and *Tbg*ISG64 were recombinantly produced overnight at 22°C in *E. coli* T7 shuffle cells (New England Biolabs) after induction of protein expression by addition of isopropyl-β-D-thiogalacto-pyranoside (IPTG) to a final concentration of 1 mM. Cells were harvested by centrifugation for 15 min at 6.000xg, 4°C. Cell pellets were resuspended in Buffer A (20 mM Tris, 500 mM NaCl, 10 mM imidazole, pH 8.0), to which phenylmethylsulfonyl fluoride (PMSF) was added at a final concentration of 1 mM immediately before cell lysis. Cells were lysed using an EmulsiFlex-C3 (AVESTIN Europe) with 1000 to 1100 bar lysis pressure at 4°C. Cell debris were removed by centrifugation for 45 min at 20.000xg, 4°C. Soluble protein was purified using immobilized-metal affinity chromatography (IMAC) by application of the cleared supernatant to nickel-nitrilotriacetic acid (NTA) beads (Qiagen) in a gravity flow column (Bio-Rad), pre-equilibrated with Buffer A. The resin was washed with 10 column volumes (CV) Buffer A twice before eluting the bound protein fraction with 10 CV Buffer B (20 mM Tris, 500 mM NaCl, 400 mM imidazole, pH 8.0). The eluate was fractionated, and those fractions containing the protein of interest were identified via SDS-PAGE, pooled, transferred into SnakeSkin Dialysis tubing (Thermo Fisher Scientific) and dialyzed overnight into Buffer C (20 mM Tris, 500 mM NaCl, pH 8.0) at 4°C. The dialyzed eluate was concentrated using Amicon Ultra centrifugal filters (Merck Millipore) with a 10 kDa MWCO before being subjected to size exclusion chromatography (SEC) on a Superdex 200 16/60 column (GE Healthcare) connected to an Äkta FPLC system (GE Healthcare), pre-equilibrated with Buffer D (20mM HEPES, 150 mM NaCl, pH 7.5). 1.5 mL elution fractions were collected throughout the run, fractions containing the protein of interest, as determined by SDS-PAGE, were pooled, flash-frozen in liquid-nitrogen and stored at -80°C until further use.

*T. b. brucei and T. b. gambiense VSGs. T. b. brucei* LiTat1.5 and *T. b. gambiense* LiTat3.1 (*Tbb*VSG LiTat1.5 and *Tbg*VSG LiTat3.1, respectively) were obtained from native source as described in [30] and [18], respectively.

### Hydrogen-deuterium exchange mass spectrometry

Hydrogen deuterium exchange was initiated by 10-fold dilution of *Tbg*ISG43, *Tbg*ISG65 or *Tbg*ISG75 into deuterated buffer (20 mM HEPES, 150 mM NaCl, pD 7.5). 50 μL aliquots (100 pmols) were taken after 20 s, 120 s and 1200 s of incubation in deuterated buffer and quenched by the addition of 50 μL of 4 M urea, 1 M glycine and 200 mM TCEP at pD 2.3, followed by immediate flash freezing in liquid nitrogen. Aliquots were quickly thawed and injected onto a Nepenthesin-2/pepsin column (AffiPro, Czech Republic). Generated peptides were trapped and desalted via a Micro-trap column (Luna Omega 5 μm Polar C18 100 Å Micro Trap, 20 x 0.3 mm) for 3 min at a flow rate of 200 μL/min using an isocratic pump delivering 0.4% (v/v) formic acid in water. Both the protease and the trap column were placed in an icebox. After 3 min, peptides were separated on a C18 reversed phase column (Luna® Omega 1.6 μm Polar C18 100 Å, 100 x 1.0 mm) and analyzed using a timsToF Pro mass spectrometer (Bruker Daltonics, Billerica, MA). Peptides were separated by a linear gradient of 10 - 30% B over 18 min, where solvent A was 2% (v/v) acetonitrile/0.4% (v/v) formic acid in water and solvent B was 95% (v/v) acetonitrile/4.5% (v/v) water/0.4% (v/v) formic acid. The mass spectrometer was operated in positive MS mode. Spectra of partially deuterated peptides were processed by Data Analysis 4.2 (Bruker Daltonics, Billerica, MA) and by in-house program DeutEx [31].

### Small angle X-ray scattering and ensemble modelling

All experiments were performed at the BioSAXS beamlines SWING (SOLEIL, Gif-sur-Yvette, France, [32]) and BM29 (ESRF, Grenoble, France, [33]). SEC-SAXS data were collected in HPLC mode using a Shodex KW404-4F column pre-equilibrated with SAXS buffer. Samples were concentrated on site to ∼10 mg/mL using a 10-kDa cutoff centrifugal filter (Amicon). Eighty-microliter samples were injected and eluted at a flow rate of 0.2 mL/min, while scattering data were collected with an exposure time of 750 msec and a dead time of 10 msec. The scattering of pure water was used to calibrate the intensity to absolute units [34]. Data were processed using CHROMIXS [35] and analyzed using BioXTas RAW [36] and the ATSAS package [37]. The information on data collection and derived structural parameters is summarized in Supplemental Table S1.

Molecular models were generated with AlphaFold2 [38] (*Tbg*ISG43, *Tbg*ISG65, *Tbg*ISG75) and AlphaFold-Multimer [39] (*Tbb*VSG LiTat1.5, *Tbg*VSG LiTat3.1). Theoretical scattering curves of the AlphaFold2 models and their respective fits to the experimental data were calculated using FoXS [40]. SAXS-based ensemble modelling was carried out using BILBOMD [40–42]. For all runs, 800 conformations were generated per R_g_, with minimal and maximal R_g_ values set at 7% and 35% of the experimentally determined Rg, respectively. Molecular graphics visualization and analysis were performed with UCSF ChimeraX [43] and PyMOL Molecular Graphics System (Schrödinger).

### Circular dichroism spectroscopy

Far-UV CD experiments were carried out on a Jasco J-1500 spectropolarimeter with a 0.2 mm path cell. Proteins were dissolved in 20 mM HEPES pH 7.5, 150 mM NaF at a concentration of 0.4 mg/mL. Spectra were recorded between 195 and 260 nm wavelength at an acquisition speed of 10 nm/min and corrected for buffer absorption. During measurements, the temperature was kept constant at 25°C. The raw CD data (ellipticity θ in mdeg) were normalized for the protein concentration and for the number of residues, according to the equation below, yielding the mean residue ellipticity ([θ] in deg cm^2^ mol^−1^), where *MM, n, C*, and *l* denote the molecular mass (Da), the number of amino acids, the concentration (mg/mL), and the cuvette path length (cm), respectively.

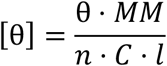

Secondary structure analysis was performed with the CD Pro software package.

### Mass spectrometry

Single gel bands were cut, chopped into small pieces, reduced with dithiothreitol, alkylated with chloroacetamide and digested with trypsin overnight. Peptides were extracted from each gel piece, lyophilized in a speed-vac, dissolved in 0.1% formic acid and 20% of the dissolved sample was separated on an UltiMate 3000 RSLCnano system (Thermo Fisher Scientific) coupled to a Orbitrap Fusion Lumos mass spectrometer (Thermo Fisher Scientific). The peptides were trapped and desalted with 2% acetonitrile in 0.1% formic acid at a flow rate of 5 μL/min on an Acclaim PepMap100 column (5 μm, 5 mm by 300 μm internal diameter (ID); Thermo Fisher Scientific). Eluted peptides were separated using an Acclaim PepMap100 analytical column (2 μm, 50 cm × 75 μm ID; Thermo Fisher Scientific). Using a constant flow rate of 300 nL/min, a 65 min elution gradient was started at 5% B (0.1% formic acid in 99.9% acetonitrile) and 95% A (0.1% formic acid). The gradient reached 30% B at 52 min, 90% B at 53 min, and was then kept constant until 57 min before being reduced to 5% B at 58 min. For the first minute, nanospray was set to 1600 V, 350 °C source temperature, measuring the scans in the range of m/z 350–2000. An orbitrap detector was used for MS with the resolution 120,000, the AGC target value was set as custom with a normalized AGC target of 250%. Maximum injection time was set to 50 ms, MSMS was acquired also using orbitrap with resolution 30,000, the data were acquired in a data-dependent manner, ions were fragmented by HCD collision energy set to 30% with dynamic exclusion set to 60 s.

The resulting raw data files were searched to identify the peptides by PEAKS (PEAKS Studio 10.0) software. PEAKS DB database search was set to search for Carbamidomethylation on Cysteine as fixed modification, Deamidation on N or Q, Oxidation on M and glycosylation specified as follows (Hex(2) HexNAc, HexNAc(3), Hex2 dHex2) was set as the variable modifications. The PEAKS DB was followed by PEAKS PTM and SPIDER.

The data files were processed also by MASCOT (Version: 2.6.2). To address the range of modifications, the error tolerant search was applied.

In all searching steps, the *T. b. gambiense* protein database (downloaded from UniProt on 31^st^ October 2022) was used.

## RESULTS

### Single-conformer AlphaFold2 models do not accurately represent the in-solution behavior of variant and invariant surface glycoproteins

To explore the conformational dynamics of ISGs and VSGs in solution, this study employed a combination of AlphaFold2-based structure prediction for generating all-atom structural models, and the collection of small angle X-ray scattering (SAXS) data for model validation. The structures of both VSGs and ISGs could be predicted with relatively high confidence as judged from local pLDDT values and overall zDOPE and pTM scores (Supplementary Figure 1). For both VSGs (*Tbb*VSG LiTat1.5 and *Tbg*VSG LiTat3.1), especially the N-terminal domains (NTDs) and the dimer interfaces are modelled with high accuracy, as evidenced by various validation metrics (PAE, pDockQ, and ipTM). Structural alignment reveals that *Tbb*VSG LiTat1.5 is a Class A1 VSG (top lobe structures reminiscent of VSG13 [44]), while *Tbg*VSG LiTat3.1 can be classified as a Class A2 VSG [45] (N-terminal lobe structure as found in VSG1, Figure 1A). In addition, the occurrence of one C-terminal domain (CTD) is predicted per monomer, albeit with lower confidence scores. Interestingly, the *Tbg*VSG LiTat3.1 AlphaFold2 model displays a relatively long C-terminal helix. While the pLDDT value analysis of this region would suggest a reliable prediction, such structures have not yet been described for VSG C-terminal regions. The structure prediction of the ISGs under study (*Tbg*ISG43, *Tbg*ISG64, *Tbg*ISG75) follows a similar trend. The N-terminal domains are modelled with high accuracy and are in accordance with the recently published, experimental structure of *Tbg*ISG65 (Figure 1A). In contrast, the ISG43 and ISG64 C-terminal domains display significantly lower pLDDT scores (<50; Supplementary Figure 1). For *Tbg*ISG75, AlphaFold2 predicts the C-terminus to consist of two long, intertwined, antiparallel α-helices with reasonably good pLDDT scores (Supplementary Figure 1).

**Figure 1.**
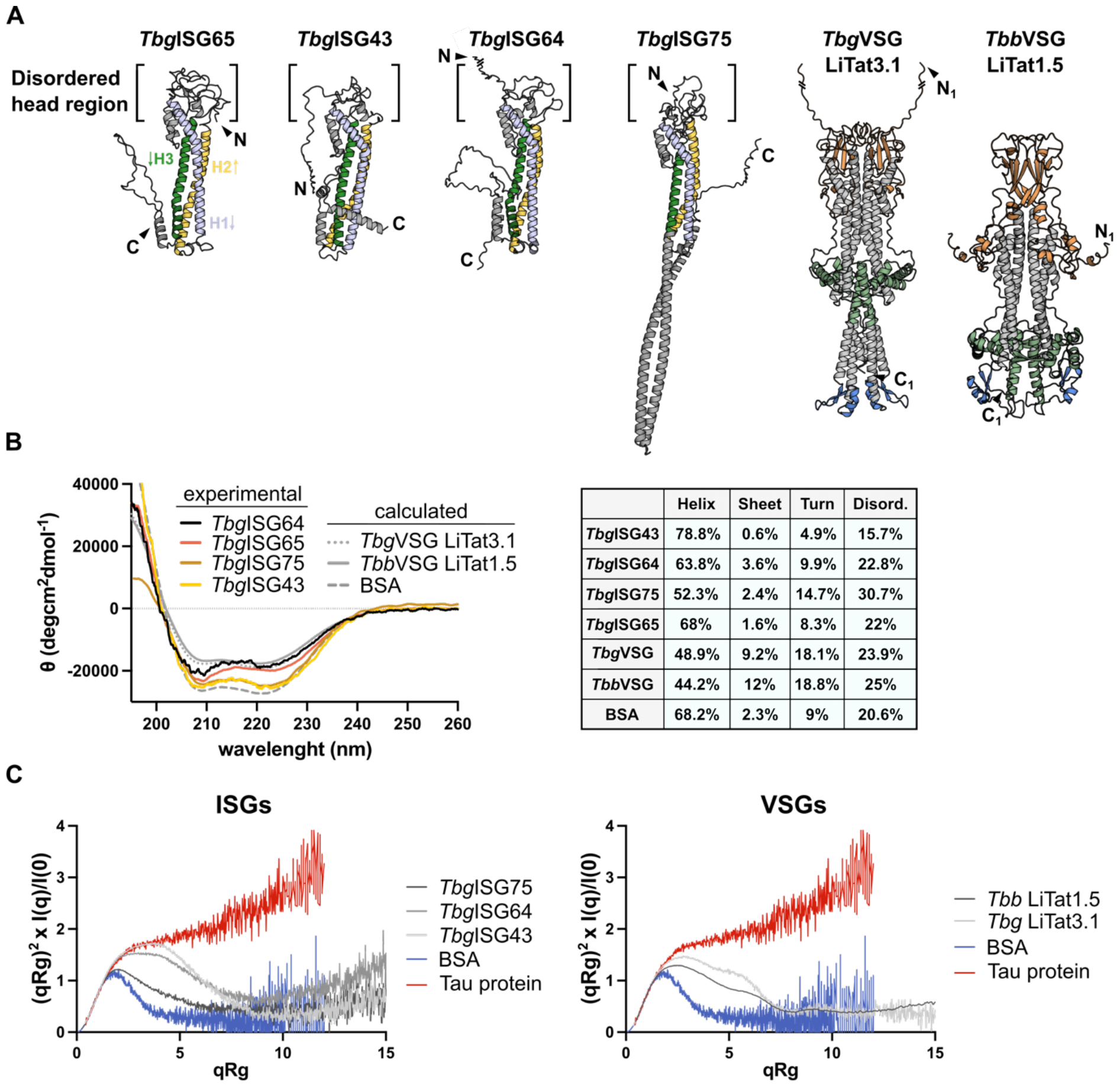
The VSG and ISG solution structures exhibit a high degree of flexibility. **(A)** The predicted AlphaFold2 models of *Tbg*ISG43, *Tbg*ISG64, *Tbg*ISG75, *Tbg*VSG LiTat3.1 and *Tbb*VSG LiTat1.5 illustrated in cartoon representation, depicted next to the hybrid structure of *Tbg*ISG65_18-363_ (PDBDEV_00000201). The three helices in the *Tbg*ISGs constituting the canonical three helix bundle have been colored uniformly and labelled on *Tbg*ISG65, arrows indicate the direction of the peptide chain from N-to C-terminus. Bottom and top lobe as well as C-terminal domains of the VSGs are highlighted in orange, green and blue, respectively. For representation purposes, the N-termini of *Tbg*ISG64 and *Tbg*VSG have been shortened, deleted residues are indicated with a line break (full length models are shown in Figure 2). **(B)** CD spectra for *Tbg*ISGs (experimental), VSGs (calculated from AlphaFold2 models) and BSA (calculated from PDB 4F5S). Secondary structure analyses (performed using CONTINLL [46,47]) for the respective spectra are summarized in the table. All CD spectra exhibit minima at 208 and 222 nm, characteristic for a largely alpha-helical fold. Similarly, all proteins have a high content of turns and disordered regions. **(C)** Dimensionless Kratky plots (R_g_) for *Tbg*ISGs (left) and *Tbb*VSG LiTat1.5 and *Tbg*VSG LiTat3.1 (right). Comparison of the plots for *Tbg* and *Tbb* proteins to reference samples for strictly folded (BSA, blue) and completely disordered proteins (Tau protein, red) demonstrate that all measured proteins contain significant fractions of both ordered and disordered components.

A first insight into the solution behavior of the studied proteins is offered by circular dichroism (CD) spectroscopy (Figure 1B). As expected, the CD spectra show a high α-helical content, but also reveal a significant amount of intrinsic disorder, which is not apparent in the single-conformer AlphaFold2 models. In both VSGs and *Tbg*ISG75, ∼50% of the residues can be found in low complexity regions (turns and intrinsic disorder). This observation is further supported by the normalized Kratky plots (indicators for intrinsic disorder), which show that the solution structures of both VSGs and ISGs are characterized by a significant degree of flexibility (Figure 1C). The latter suggests that the VSG and ISG solution structures would be best described by a conformational ensemble. Indeed, as shown in Figure 2, single-conformer AlphaFold2 models are insufficient to explain the solution behavior of the VSG and ISG molecules under study. Theoretical scattering curves derived from the static models differ significantly from the experimental SAXS curves as indicated by the poor fits and high χ^2^ values. The largest disagreements between predicted and solution structures were observed for *Tbb*VSG LiTat 1.5 (χ^2^ = 522) and *Tbg*ISG75 (χ^2^ = 150).

**Figure 2.**
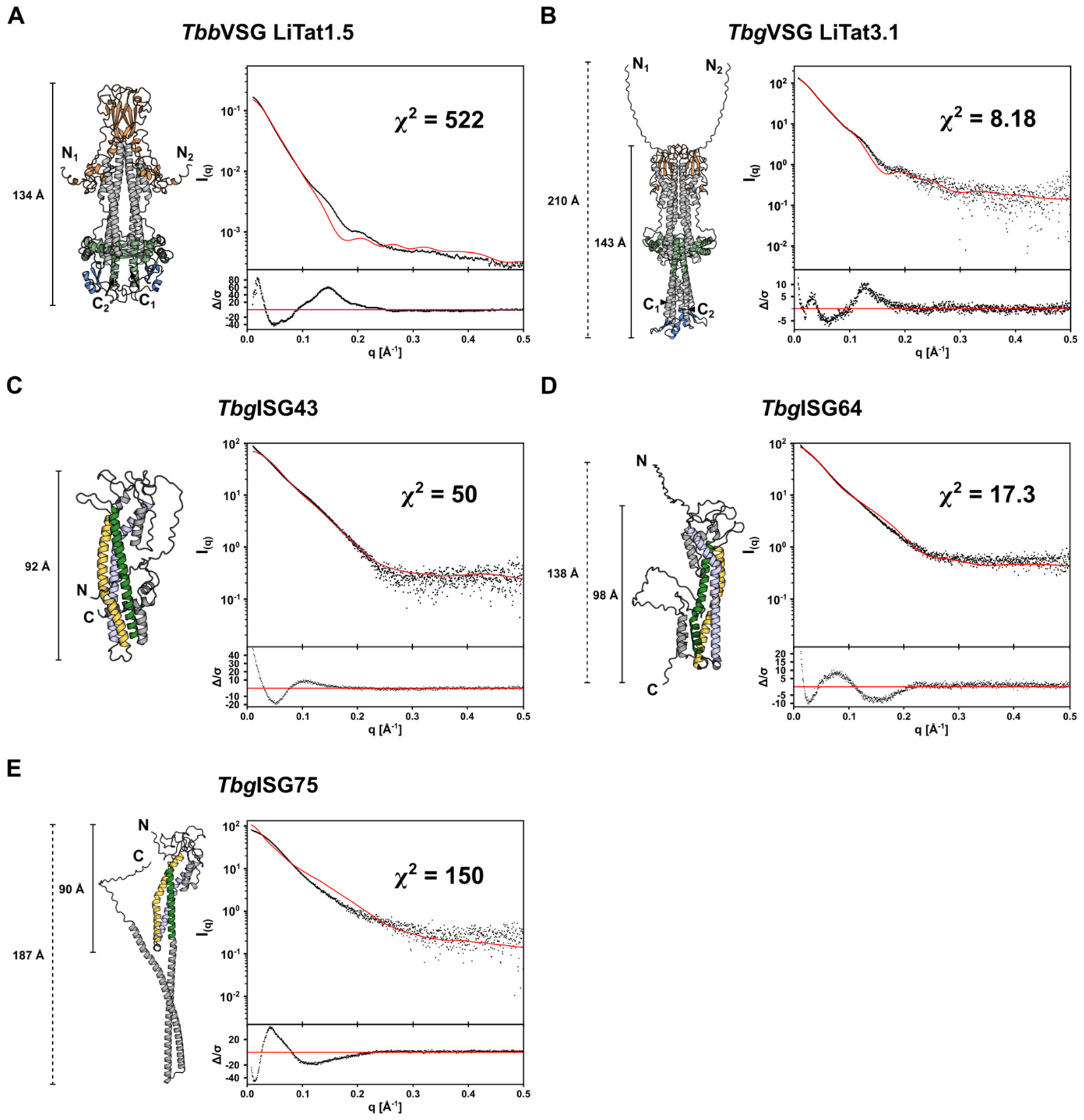
The VSG and ISG single-conformer AlphaFold2 models do not account for the experimentally observed solution behavior. The theoretical scattering curves of the unaltered AlphaFold2 models for *Tbb*VSG LiTat1.5 (A), *Tbg*VSG LiTat3.1 (B), *Tbg*ISG43 (C), *Tbg*ISG64 (D) and *Tbg*ISG75 (E) were calculated (red curves) and compared to the experimental SAXS data (black dots). The fit residuals are displayed as insets below the scattering curves. The respective AlphaFold2 models used for the calculations are shown (colored using the same scheme as in Figure 1).

### VSGs adopt distinct structural states in solution that are best described by conformational ensembles

Structural flexibility of VSGs (*Tbb*VSG MiTat1.1 and *Tbb*VSG IlTat1.24) has previously been investigated [18]. Bartossek and co-workers employed a SAXS-based, rigid body modelling approach to demonstrate that the VSG solution behavior is best described by a conformational ensemble, in which the NTDs behave as rigid bodies, while the CTDs are flexible. In line with these findings, we found that the static AlphaFold2 models of *Tbb*VSG LiTat1.5 and *Tbg*VSG LiTat3.1 inadequately describe the solution behavior (Figure 2, panels A and B).

By employing an alternative, molecular dynamics-based method for generating and selecting conformers during SAXS-driven ensemble modeling, we could confirm the initial findings of Bartossek et al. As shown in Figure 3A, the CTD of VSGs is highly flexible, thereby allowing the molecule to adopt more than one conformational state. For *Tbb*VSG LiTat1.5, the experimental scattering data was best explained by an ensemble (χ^2^ = 2.00), 40% of which consists of a compact conformer (R_g_ = 42.16 Å, D_max_ = 155 Å), while the remaining 60% comprises conformers with varying degrees of extension (R_g_ = 47.6 Å, D_max_ = 196 Å for the most extended conformer). The application of the same protocol to *Tbg*VSG LiTat3.1 reveals that the best fit to the experimental data is provided by a conformational ensemble in which the CTD is also granted full conformational flexibility (χ^2^ = 1.75, Figure 3B), contrasting the AlphaFold2 model discussed above (Figure 1A). The majority of the ensemble (78.5%) comprises a compact conformation (R_g_ = 43.57 Å, D_max_ = 167 Å), while the remaining 21.5% corresponds to an extended conformation (R_g_ = 48.67 Å, D_max_ = 211 Å).

**Figure 3.**
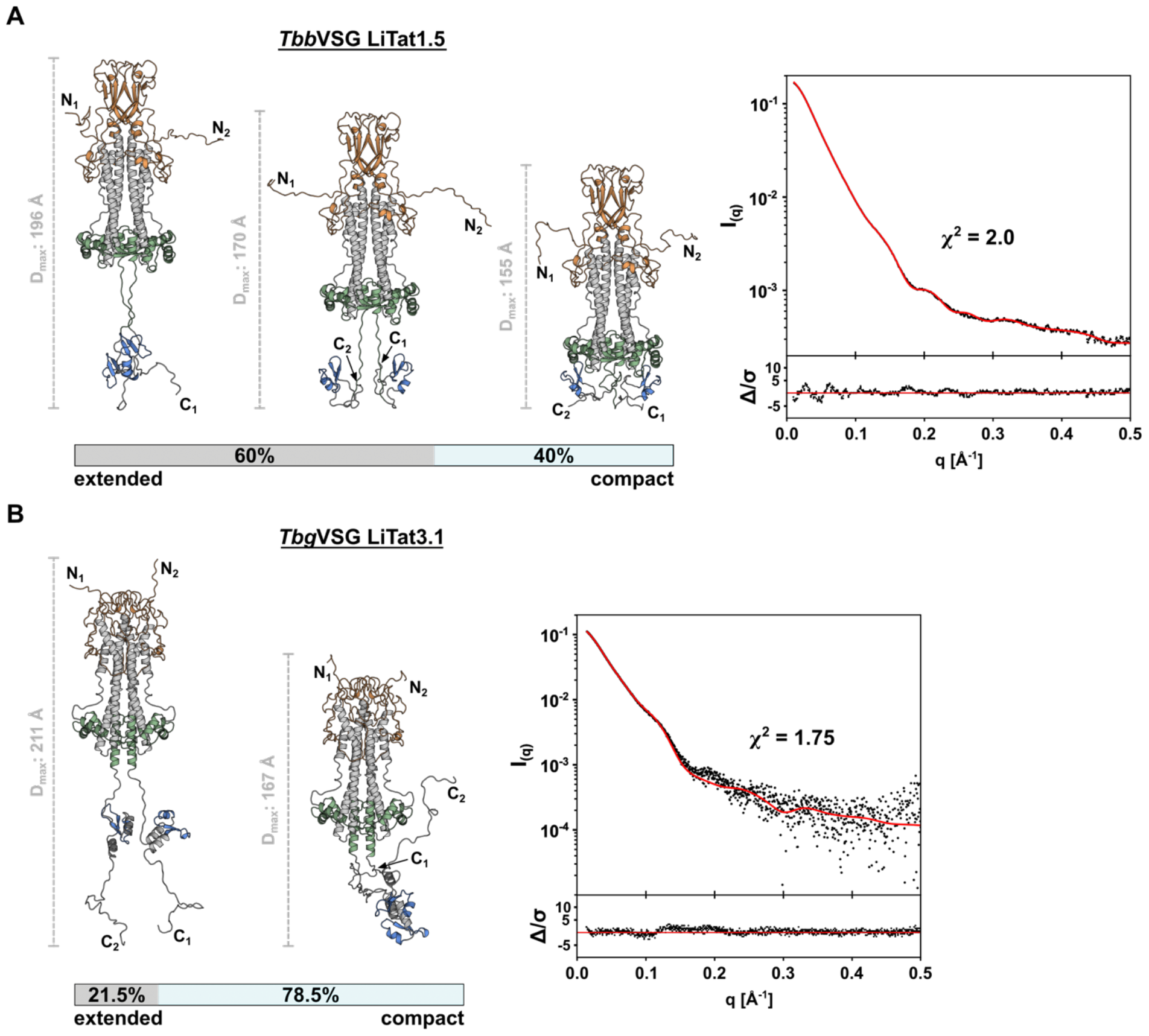
Conformational ensemble modelling reveals structural flexibility of VSGs in solution. **(A)** The solution scattering of *Tbb*VSG LiTat1.5 is best described by a conformational ensemble composed of 5 models. Three representative conformations are shown, alongside a scale bar indicating the D_max_ of the individual conformer. **(B)** The SAXS data of *Tbg*VSG LiTat3.1 is best described by a conformational ensemble composed of 2 models. Both conformations are depicted next to a scale bar indicating the D_max_ of the individual conformer. In both panels, the experimental scattering data (black dots) and calculated ensemble scattering curves (red line) are shown. The residuals of the fit are shown below.

In contrast to the relatively small differences in D_max_ (up to 16 Å) between the 2 conformational states of *Tbb*VSGs reported by Bartossek et al., the differences in D_max_ found for the compact and the most extended conformers of *Tbb*VSG LiTat1.5 and *Tbg*VSG LiTat3.1 are much larger with 41 Å (Figure 3A) and 44 Å (Figure 3B), respectively. Furthermore, the most extended conformers of the VSGs investigated here exceed the D_max_ reported for VSGs by Bartossek et al. by approximately 50 Å.

### The membrane-proximal, C-terminal regions of ISGs are intrinsically disordered

Considering that low pLDDT scores (<50) can serve as an indicator for disorder [48], the AlphaFold2 models presented in this study suggest the presence of intrinsic disorder in the C-terminal regions of *Tbg*ISG43 and *Tbg*ISG64. To investigate this further, we employed hydrogen-deuterium exchange mass spectrometry (HDX-MS) (Figure 4). Differences in relative deuteration rates can provide insight into protein secondary structure, as protein regions with low complexity typically exhibit faster deuteration rates than their well-folded counterparts [49]. This is clearly illustrated by *Tbg*ISG65, which serves as a benchmark here, demonstrating that deuteration rates exhibit a strong correlation with the experimental structure (Figure 4, bottom panel). Similarly, for *Tbg*ISG43, *Tbg*ISG64 and *Tbg*ISG75, residues predicted by AlphaFold2 with high confidence to constitute the canonical three-helical bundle show significantly lower deuteration rates compared to the rest of the protein. Underlining the low pLDDT scores, the HDX-MS analysis also confirms a disordered N-terminus for all proteins analyzed, as indicated by the fast deuteration rates. Notably, *Tbg*ISG43 N-terminal residues Gly2-Leu14 appear to exhibit a lower deuteration rate than residues Val25-Glu39, indicating a stable secondary structure. This however is most likely an artifact caused by technical limitations, such as insufficient proteolytic digest or low ionization efficiency. Gaps in the sequence coverage, caused by the lack of a sufficient quantity of smaller peptides from that part of the protein, may result in a loss of resolution, producing nonrepresentative deuteration rates (Supplementary Figure 2). In all proteins analyzed, the residues predicted to form membrane-distal, disordered head domains undergo a large degree of deuteration already after 20 s. The C-terminal residues of *Tbg*ISG43, *Tbg*ISG64 and *Tbg*ISG65 (predicted to be largely disordered) deuterate to a similar degree as the disordered loops in the head domains, supporting the AlphaFold2 prediction. C-terminal residues *Tbg*ISG43 Val328-Arg348 are predicted to form an α-helix, which appears to be supported by the slow initial deuteration rate over the first 20 s. The same observation can be made for *Tbg*ISG64 residues Ala354-Leu360.

**Figure 4.**
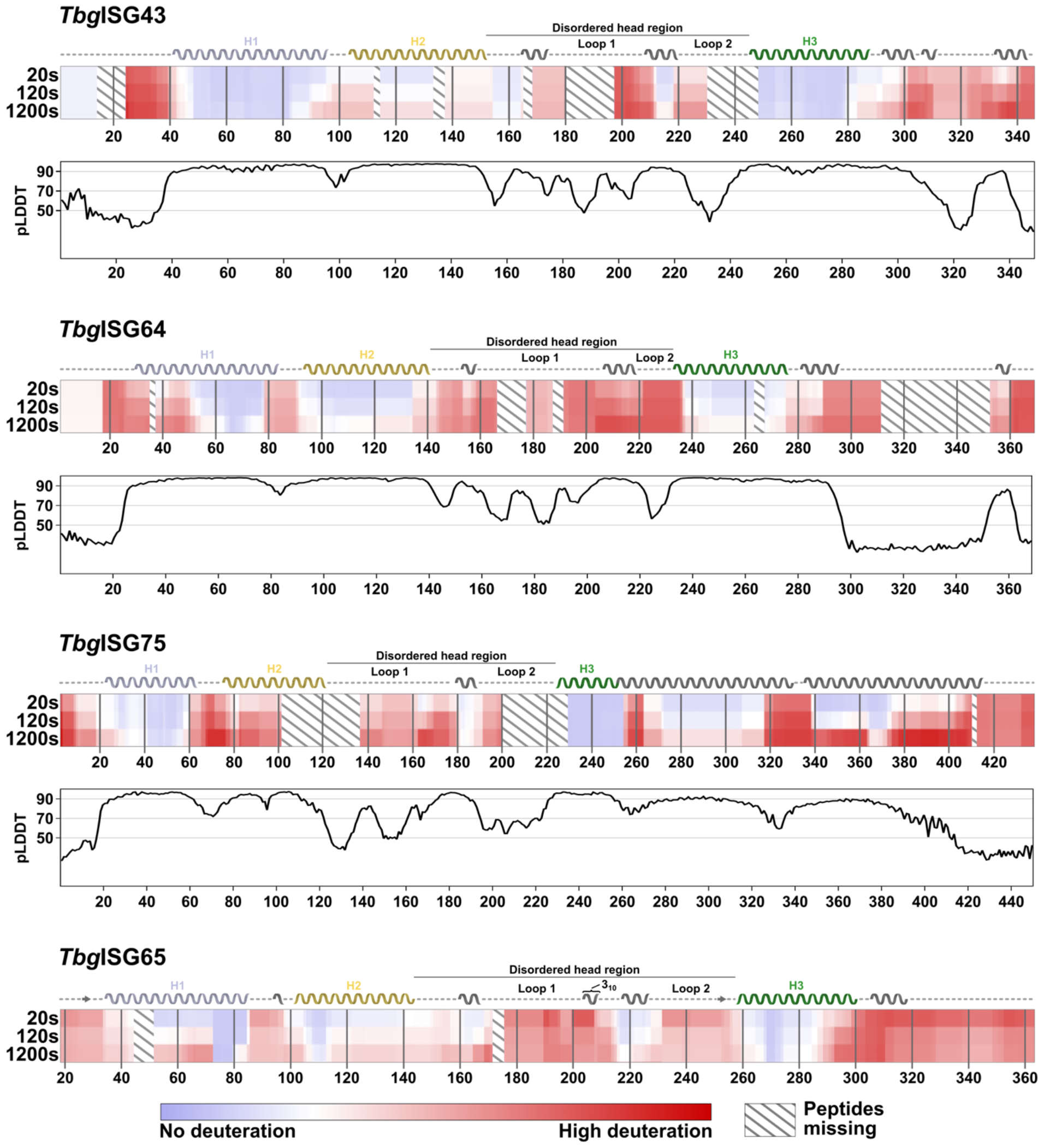
HDX-MS reveals intrinsically disordered regions in TbgISGs. Chiclet plots showing the relative deuteration across the sequences of *Tbg*ISGs at 20, 120 and 1200 s of incubation. The amount of relative deuteration is indicated by color gradients, ranging from blue (no deuteration) to red (high deuteration). Peptides absent in the analysis are represented by diagonal lines. For *Tbg*ISG75, *Tbg*ISG64 and *Tbg*ISG43, the secondary structure, as predicted by AlphaFold2, is indicated above the corresponding chiclet plot. For *Tbg*ISG65, the secondary structure of the integrative *Tbg*ISG65 hybrid structure deposited to the PDBDev (PDBDEV_00000201) is shown [7]. Disordered regions are represented by a dashed line, beta sheets as arrows and alpha helices as helices. The three helices constituting the canonical three-helical bundle are labelled (H1 to H3) and colored as in previous figures. The 3_10_-helix in ISG65 is indicated with a label. The disordered loops forming the disordered, membrane-distal head region of the ISGs are labeled accordingly. Underneath the chiclet plots of *Tbg*ISG43, *Tbg*ISG64 and *Tbg*ISG75, the per-residue pLDDT scores of the respective AlphaFold2 models are plotted. Thresholds for high (>90), medium (>70) and low (<50) modelling confidence are indicated with grey lines.

HDX-MS analysis of *Tbg*ISG75 however contradicts the AlphaFold2 model (Figure 1C) as this experimental evidence does not support the existence of the two predicted, continuous C-terminal helices in solution (residues Lys262-Ala330 and Glu336-Gly417). Unlike for the other *Tbg*ISGs studied, the HDX-MS analysis furthermore suggests that the C-terminus of *Tbg*ISG75 is not entirely disordered either. Residues Arg272-Arg317 initially exhibit significantly lower deuteration rates than truly disordered parts of the protein, such as the head domains, but show increased deuteration after longer incubation periods when compared to the predicted helices of the canonical three-helical bundle. A similar trend can be observed for residues Lys339-Ala370, and to a lesser degree for residues Glu371-Glu408. After incubation for 1200 s, residues Arg317-Glu438 show a large degree of deuteration, suggesting a high degree of flexibility. This indicates that parts of the C-terminus of *Tbg*ISG75 undergo structural transitions and therefore may occur as both folded and disordered in solution.

In line with a high χ^2^ (and hence a disagreement between the *Tbg*ISG75 Alphafold2 and solution structures), HDX profiles also indicate that the prediction of an unusually long helix 3 is incorrect. High deuterium incorporation shown for residues Lys255-Ser264 strongly suggest that helix 3 terminates around residue Lys255.

Hence, in conclusion, these data show that the membrane-proximal, C-terminal regions of *Tbg*ISG43, *Tbg*ISG64, and *Tbg*ISG75 are characterized by a significant degree of intrinsic disorder. This discovery aligns with the previously described observation that, similarly to the VSGs, the single-conformer AlphaFold2 models for the *Tbg*ISGs proved inadequate for explaining the solution scattering data (Figure 2, panels C to E).

### Intrinsic disorder enables T. b. gambiense ISGs to switch between compact and elongated conformational states

To improve the fit to the experimental data and, thus, provide structural models that represent the *Tbg*ISG solution behavior best, the HDX-MS data were employed as restraints in the SAXS-based ensemble modelling (Supplementary Table 1). In all three cases, the best results were obtained when the structures are described by a conformational ensemble, in which the membrane-distal, N-terminal domains are treated as rigid bodies and the membrane-proximal, C-terminal regions are disordered. A thorough investigation of the conformational properties revealed that all ensembles yielding a good fit to the experimental data consist of two conformers: a compact and a highly extended conformer (Figure 5).

**Figure 5.**
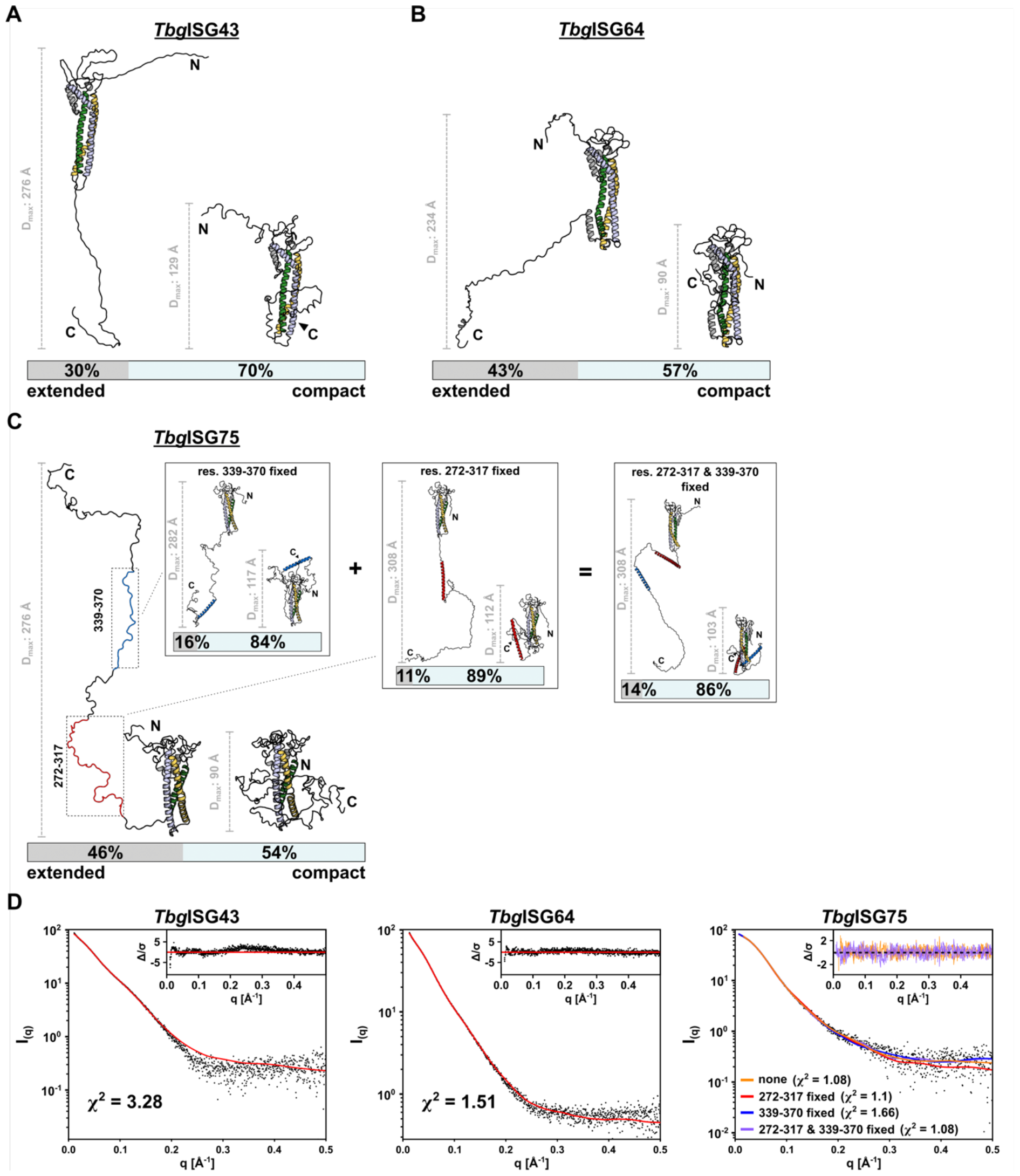
Conformational ensemble modelling reveals structural flexibility of ISGs in solution. Representative ensembles of (A) *Tbg*ISG43, (B) *Tbg*ISG64 and (C) *Tbg*ISG75 for the experimental SAXS data, calculated using BILBOMD. The smallest ensemble producing a reasonable improvement in goodness of fit of the theoretical scattering curve to the experimental data was chosen. Models constituting the selected ensemble are shown with their respective weight. For each model, the D_max_ is displayed with a representative scalebar. For ISG75 in (C) four ensembles are shown, corresponding to ensemble calculation without HDX-MS restraints, restraints applied to residues Arg272-Arg317 (highlighted in red) [left insert], residues Lys339-Ala370 (highlighted in blue) [middle insert] and both [right insert]. (D) Experimental scattering data are shown, overlaid with the theoretical scattering curves of the selected ensembles and the respective χ^2^ of the fit. Residuals are shown as insets. For ISG75, the residuals for the ensemble without restraints for modelling of the CTD (orange) and with restraints in both res. 272-317 and res. 339-370 (purple) are shown.

For *Tbg*ISG43, the selected ensemble provided a χ^2^ = 3.28, indicating a reasonable fit of the ensemble’s theoretical scattering curve to the experimental data. While 70% of the ensemble consists of a compact conformer (R_g_ = 33.47 Å, D_max_ = 129 Å), the remaining 30% corresponds to a highly elongated form (R_g_ = 69.03 Å, D_max_ = 276 Å) (Figure 5A). A similar result was obtained for *Tbg*ISG64, for which the selected two-model ensemble (comprising 57% and 43% compact and elongated conformers, respectively) could be fitted to the experimental data with a χ^2^ = 1.51.

For *Tbg*ISG75, our modelling attempts were more extensive given that the HDX-MS data indicated that the C-terminal region displays an intriguing structural promiscuity, possibly transitioning between helical and disordered forms. When treating the entire C-terminus of *Tbg*ISG75 as disordered, the best fitting ensemble (χ^2^ = 1.08) consisted of 54% compact (R_g_ = 28.37 Å, D_max_ = 90 Å) and 46% extended (R_g_ = 89.96 Å, D_max_ = 323 Å) conformers. To explore ensembles representing the HDX-MS data more closely, ensembles with α-helical elements in their C-terminal regions corresponding to lower deuteration rates were generated. Among these ensembles, the least optimal fit between theoretical and experimental scattering data was obtained when restraining conformational sampling in residues Arg272-Arg317 (χ^2^ = 1.66), conserving the predicted α-helical fold for these residues. Even though the distances spanned by the compact (R_g_ = 31.78 Å, D_max_ = 112 Å) and the extended (R_g_ = 95.2 Å, D_max_ = 308 Å) conformers did not change dramatically, the fraction of conformers in each state did, with only 11% of the molecules modelled in the extended conformation. When excluding residues Lys339-Ala370 from the conformational sampling, thereby enforcing the predicted α-helical secondary structure in this region, a comparable distribution was observed. Although the fractions of the two conformers contributing to the ensemble did not change much with 16% of elongated and 84% of compact conformers, the calculated distances the two conformations spanned were noticeably different (D_max, compact_ = 117 Å and D_max, extended_ = 282 Å). This ensemble appears to describe the experimental data better in comparison with the previous restrains as the χ^2^ improved to 1.1. Finally, when applying α-helix restrains in both regions of the C-terminus simultaneously, an ensemble with a similar distribution, 14% compact (R_g_ = 31.07 Å, D_max_ = 103 Å) and 86% extended (R_g_ = 88.79 Å, D_max_ = 308 Å) conformers was produced. Interestingly, the theoretical scattering could be fitted to the experimental data with a χ^2^ = 1.08, identical to the ensemble with no restraints in the C-terminus. These findings correlate well with the HDX-MS data, supporting the hypothesis that both states, entirely disordered and partially folded C-termini, may be present in solution. Noticeably, the presence of one or two helices in the CTD shifted the equilibrium from an equal distribution of compact and extended conformations, as observed in a fully disordered CTD, towards a prevalence of compact states.

## DISCUSSION

The advent of AI-based structure prediction programs such as AlphaFold2 [38] and RoseTTAFold [50] has revolutionized and democratized structural biology. These tools can readily provide highly accurate, all-atom structural models without any experimental input. However, the generated structures do not always capture the solution behavior of proteins and their complexes, thus providing a very limited insight into their potential biological functions. This is especially true for intrinsically disordered proteins (IDPs) or proteins containing intrinsically disordered regions (IDRs). In these cases, AlphaFold2 structural models can be employed in conjunction with SAXS-based conformational sampling to provide an accurate description of a protein’s conformational ensemble in solution [51,52]. Such an approach was applied here to investigate the structural features of *Tbg*ISGs (*Tbg*ISG43, *Tbg*ISG64, and *Tbg*ISG75), interpret the results within the context of the *Tbg*VSG layer and provide a comparison with recent work published on *Tbg*ISG65 [7].

We first sought to validate the SAXS-based modeling approach employed here by collecting solution scattering data for *Tbb*VSG LiTat1.5 and *Tbg*VSG LiTat3.1 and comparing these to the results previously reported for *Tbb*VSG MiTat1.1 and *Tbb*VSG IlTat1.24 [18]. In their work, Bartossek et al. showed that the *Tbb*VSG umbrella extends approximately 140 Å to 155 Å from the parasite surface. For both VSGs employed in this study, the solution behavior is best described by conformational ensembles with average D_max_ values of approximately 170 Å. Considering that in both studies the proteins were directly obtained from the parasite plasma membrane by GPI-specific phospholipase C cleavage, thereby eliminating all steric restraints imposed by localization on the parasite’s membrane (e.g. the presence of neighboring molecules and protomer GPI-anchoring), we found our solution scattering data and resulting ensemble models to be in good agreement with the published *Tbb*VSG results [18]. Notably, the observed maximum dimensions of the extended models within the ensembles were larger in our approach than previously reported. It is conceivable that an MD-based approach can sample a larger conformational space of intrinsic disorder than the rigid body modelling based protocol employed earlier. Additionally, although the VSGs used in these studies belong to the same protein family, they are not identical, thus potentially explaining the observed differences.

After this validation step, the same SAXS-based modeling approach was employed to investigate the solution behavior of *Tbg*ISG43, *Tbg*ISG64, and *Tbg*ISG75. For all three *Tbg*ISGs, we identified a clear requirement for ensemble-based structural modelling, *i*.*e*., while single-conformer AlphaFold2 models could not adequately explain the SAXS data, conformational ensembles accurately describe the solution scattering data. Our findings show that the *Tbg*ISGs possess a C-terminal, membrane-proximal IDR, constituting a flexible tether between the TMD and the well-structured NTD. This configuration allows exploration of a large conformational landscape, encompassing highly compact to maximally extended conformers. For most ISGs the intrinsic disorder of these tails was captured well by AlphaFold2 (these regions have pLDDT scores around 50, which is indicative of intrinsic disorder [48]). However, the C-terminus of *Tbg*ISG75 was predicted to form a long pair of coiled α-helices, with confidence scores ranging from high (pLDDT between 70 to 90) to very high (pLDDT ≥ 90). Interestingly, a recent study has found AlphaFold2 capable of identifying conditionally folded IDRs and IDPs [53]. Alderson et al. have reported that secondary structure predictions with high and very high confidence scores in protein regions identified as IDRs via sequence-based prediction software are likely to resemble conditionally folded conformations, induced either upon post-translational modification or ligand binding. Coincidentally, most conditionally folded IDRs predicted by AlphaFold2 are helical. Considering the long-suspected role of ISG75 as an abundant receptor for a yet to be discovered ligand, the apparent discrepancy between the experimentally determined conformational ensemble and the single-conformer AlphaFold2 model could suggest that the *Tbg*ISG75 C-terminal linkers are conditionally folding IDRs. Likewise, the *Tbg*VSG LiTat3.1 AlphaFold2 model contains a predicted C-terminal α-helix (Val360-Ala410) harboring high to very high pLDDT scores, which is not in accordance with its solution behavior and may potentially highlight this stretch as a conditionally folding IDR. While the molecular cue for the conditional folding remains speculative, both ligand-binding induced changes and folding in proximity of the parasite membrane are conceivable. In the absence of a stabilizing entity, the integrative structural biology approach adopted here indicates that the *Tbg*ISG75 and *Tbg*VSG LiTat3.1 solution structures are best described by conformational ensembles. For *Tbg*ISG75, the latter consists of a mixture of compact and extended conformers with transiently formed α-helices (Arg272-Arg317 and Lys339-Ala370).

The solution behavior observed for *Tbg*ISG43, *Tbg*ISG64, and *Tbg*ISG75 is reminiscent of similar findings recently published for *Tbg*ISG65 [7]. Like the *Tbg*ISGs used in this study, *Tbg*ISG65 possesses a C-terminal, disordered linker, enabling the extension beyond the boundaries of the VSG layer and thereby facilitating interaction with its ligands, human complement factor C3 and its proteolytically activated fragments [6,7,24]. In a similar fashion, the IDRs of *Tbg*ISG43, *Tbg*ISG64, and *Tbg*ISG75 enable these proteins to adopt distinct conformational states, which would allow them to either protrude from or reside within the VSG coat (Figure 6). Indeed, with compact conformers spanning 90 Å (*Tbg*ISG64, *Tbg*ISG75) to 129 Å (*Tbg*ISG43), all three *Tbg*ISGs studied here appear to be deeply embedded within the VSG coat (Figure 6A). However, it is important to note that the experimentally determined dimensions of the most compact *Tbg*ISG conformers presented here are likely underestimated. This is because the absence of steric constraints imposed by TMD-anchoring would permit the C-terminal linker to adopt conformations that are not possible within the confines of the membrane context. Nonetheless, in their compact conformation, much of the *Tbg*ISG N-terminal domain would remain buried and be located at a similar distance from the parasite surface as the VSG’s C-terminal domain (∼70 Å). Given that the latter was postulated to be difficult to target by the host immune system [54,55], such a compact molecular surface organization would effectively shield the *Tbg*ISGs from an immune response, perhaps only leaving their membrane-distal head domains exposed, protruding into the gap between adjacent VSG dimers. Using HDX-MS disorder mapping we could demonstrate that, in addition to *Tbg*ISG65, other members of the *Tbg*ISG super-family also possess these domains. This is a new feature of trypanosomal invariant surface proteins that was so far only described for VSGs where similar structures act as immune-dominant decoys [56–58]. This is in line with previous studies showing that ISG65 is generally inaccessible to antibodies in *T. brucei*, while only minimal antibody binding to ISG75 is observed in fixed but not live cells [22]. In combination with the well-known rapid endocytosis and efficient turnover of surface antigens exhibited by trypanosomes [59], the IDR-mediated retractability of ISGs to reduce surface exposure could indeed be regarded as a highly efficient evasion mechanism for invariant determinants. In contrast, the elongated *Tbg*ISG43 and *Tbg*ISG75 conformers would be capable of reaching beyond the VSG layer boundary, even when compared to the overextended solution state observed for *Tbg*VSG Litat3.1 (Figure 6B). Although *Tbg*ISG64 appears to span the smallest maximum distance of the three *Tbg*ISGs in its extended conformation (D_max_ = 234 Å), the N-terminal domain may still protrude well beyond the CTD of the VSG coat and, therefore, gain access to potential ligands. Although we believe our data to accurately depict the solution behavior of soluble ISGs and VSGs, it is possible that TMDs and GPI anchors as well as steric hindrances imposed by neighboring molecules add additional restraints on the trypanosome surface that could not be incorporated into our analysis. It also remains to be elucidated whether molecular triggers exist that can initiate vertical movement of surface receptors via their C-terminal IDRs and how ramping of variant and invariant proteins is coordinated. Our observation that the conditional presence of more secondary structure in the *Tbg*ISG75 CTD (i.e. fully disordered vs formation of 1 or 2 α-helices) yields predominantly compact conformations may provide a clue to how this could possibly be achieved.

**Figure 6.**
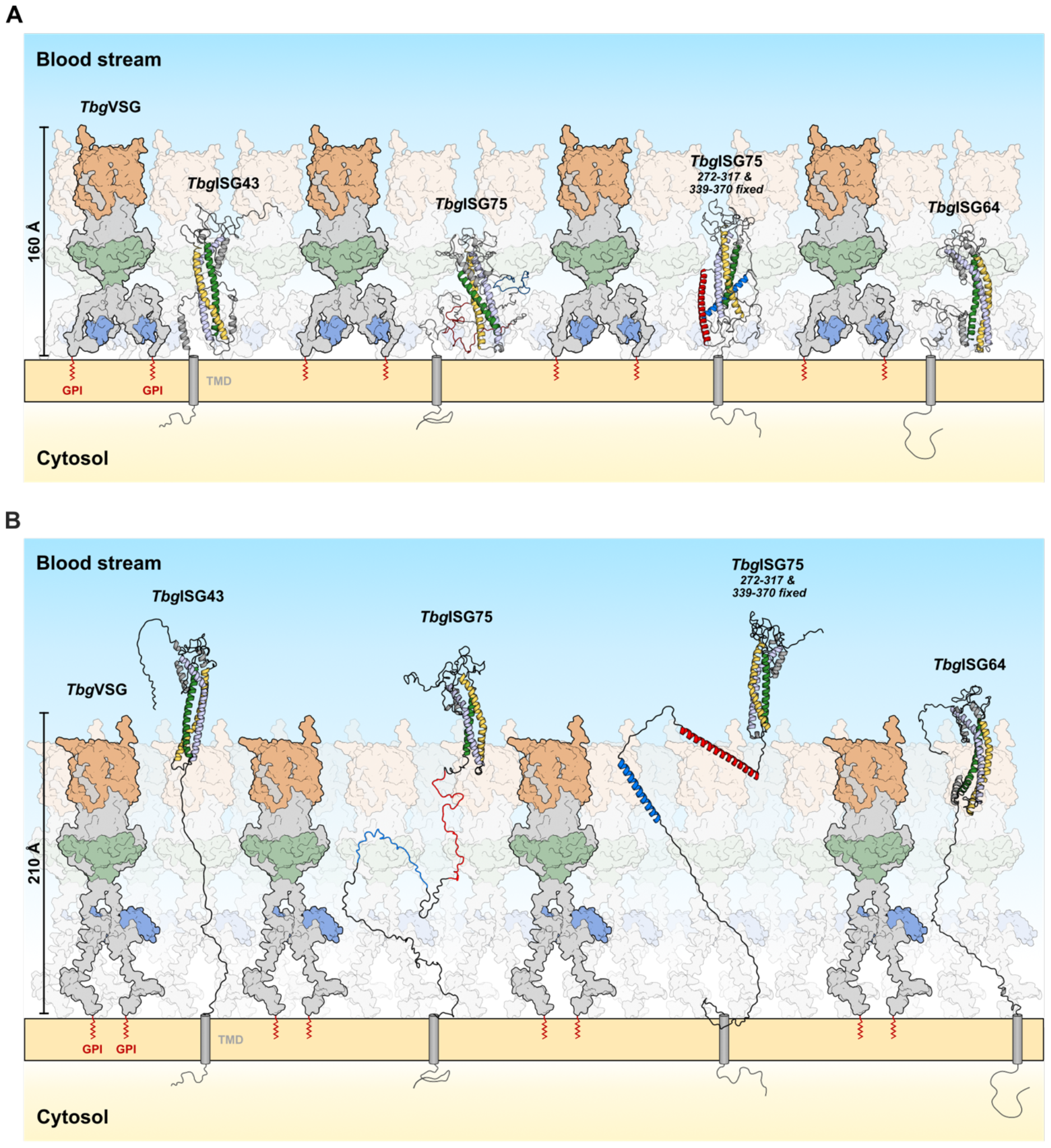
An updated model of the *Trypanosoma* surface coat – Functional implications of ISG conformational flexibility within the VSG umbrella model. **(A)** In the retracted form, ISGs reside within the VSG coat (represented here by *Tbg*VSG LiTat3.1), leaving only their disordered, loop-rich head domains accessible to molecules in the host’s blood stream, while concealing all other epitopes under the protective VSG umbrella. **(B)** In the extended conformation, both ISG43 and ISG75 protrude well beyond the boundaries of the VSG layer, even when the latter is in its extended conformation. Whilst in this model ISG64 would not be able to fully extend beyond the maximally extended VSG umbrella, it still protrudes beyond the VSGs CTD, thereby implying accessibility to potential ligands. The GPI anchors of the VSGs are depicted as a red “zigzag”, while transmembrane domains of ISGs are shown as gray cylinders. The N- and C-termini of the protein models obtained from SAXS modeling were manually modified for this figure to allow for membrane anchoring via the C-terminal linker without altering the D_max_ of the models.

The results presented in this study further expand our understanding of the structure-function relationship of *Trypanosoma* ISGs, which have been implicated in a wide range of functions (including nutrient uptake, adhesion, environmental sensing, and immune evasion). In agreement with previous reports [7,28], these findings add to a growing body of evidence observed for non-VSG surface antigens in which intrinsic disorder appears to play a prominent, functional role at the host-parasite interface. This enables us to contribute to an updated model of the trypanosome surface coat and its embedded proteins, portraying it as a highly dynamic structure rather than a static barrier.

## ACKNOWLEDGEMENTS

Research in the laboratory of S. Zoll’s lab is supported by the Czech Science Foundation (project 22-21612 S). H. Sülzen was supported by the Grant Agency of Charles University (project no. 383821/2600). R. Geens is a doctoral fellow supported by a DOCPRO4-NIEUWZAP (code 40043) grant awarded to Y.G.-J. Sterckx by the University of Antwerp ‘Bijzonder Onderzoeksfonds (BOF)’. B. Stijlemans was funded by the Strategic Research Program (SRP#47, VUB). We acknowledge the Structural mass spectrometry core facility of CIISB, Instruct-CZ Centre, supported by MEYS CR (LM2023042) and European Regional Development Fund-Project "UP CIISB” (No. CZ.02.1.01/0.0/0.0/18_046/0015974) with regard to HDX-MS measurements carried out by Petr Pompach. We extend our gratitude to Lucie Bednarová and Martin Hubalek of the IOCB for performing the CD and MS measurements, respectively. We thank Philippe Büscher from the Institute of Tropical Medicine in Antwerp, Belgium, for generously providing the *T. b. gambiense* strain that was used in this study. The authors wish to thank the staff of the SWING beamline at SOLEIL synchrotron (Javier Perez and Thomas Bizien) and BM29 beamline at ERSF (Anton Popov) for outstanding support. The authors acknowledge use of the CalcUA and VSC supercomputing facilities for AlphaFold2 structure prediction and wish to thank the staff for outstanding support.

## CONTRIBUTIONS

Y.G.-J.S. and S.Z. conceptualized the study. H.S. and S.Z. performed recombinant protein expression and purification. F.Z. and M.Z. cultured *T. b. gambiense* and provided cell pellets for VSG extraction. H.S. purified *Tbg*VSG LiTat3.1 from native source. B.S. and S.M. purified *Tbb*VSG LiTat1.5 from native source. AlphaFold2 predictions were made by Y.G.-J.S. R.G. and Y.G.-J.S. performed the SAXS measurements. SAXS data processing and analysis was done by H.S., A.N.V., S.Z. and Y.G.-J.S. HDX-MS data analysis was performed by H.S. and S.Z. SAXS driven ensemble modelling and analysis was done by H.S. (*Tbg*ISGs) and Y.G.-J.S. (*Tbb*VSG and *Tbg*VSG). The manuscript was written by H.S., Y.G.-J.S. and SZ.

