## Supplementary information for "Beyond the VSG Layer: Exploring the Role of Intrinsic Disorder in the Invariant Surface Glycoproteins of African Trypanosomes"

**Table S1.** SAXS data collection and scattering-derived parameters for the studied proteins.

| | ***Tbb*VSG LiTat1.5** | ***Tbg*VSG LiTat3.1** | ***Tbg*ISG43** | ***Tbg*ISG64** | ***Tbg*ISG75** |
|---|---|---|---|---|---|
| **Data collection parameters** | | | | | |
| Beam line | SWING (SOLEIL) | SWING (SOLEIL) | SWING (SOLEIL) | SWING (SOLEIL) | BM29 (ESRF) |
| Wavelength (Å) | 0.99 | 0.99 | 0.99 | 0.99 | 0.99 |
| $q$ range ($Å^{-1}$)* | 0.003 - 0.493 | 0.003 - 0.493 | | | |
| Concentration (mg $ml^{-1}$) (mode) | 10.00 (SEC-SAXS) | 9.85 (SEC-SAXS) | 9.96 (SEC-SAXS) | 10.96 (SEC-SAXS) | 9.80 (SEC-SAXS) |
| Buffer conditions | 50 mM Tris, 500 mM NaCl, pH 7.5 | 20 mM HEPES, 150 mM NaCl, 3% glycerol, pH 7.5 | 20 mM HEPES, 150 mM NaCl, 3% glycerol, pH 7.5 | 20 mM HEPES, 150 mM NaCl, 3% glycerol, pH 7.5 | 20 mM HEPES, 150 mM NaCl, 3% glycerol, pH 7.5 |
| Temperature (°C) | 20 | 20 | 20 | 20 | 20 |
| **Structural parameters** | | | | | |
| $I(0)$ ($cm^{-1}$) [from Guinier] | 0.18 | $0.13\ 10^{-6}$ | $0.76\ 10^{-9}$ | $0.83\ 10^{-9}$ | 79.00 |
| $R_g$ (Å) [from Guinier] | 43.94 | 46.84 | 31.77 | 31.63 | 31.89 |
| $I(0)$ ($cm^{-1}$) [from $p(r)$] | 0.18 | $0.13\ 10^{-6}$ | $0.77\ 10^{-9}$ | $0.83\ 10^{-9}$ | 79.30 |
| $R_g$ (Å) [from $p(r)$] | 46.55 | 48.20 | 33.16 | 32.28 | 32.51 |
| $I(0)$ ($cm^{-1}$) [from GPA] | 0.17 | $0.12\ 10^{-6}$ | $0.72\ 10^{-9}$ | $0.86\ 10^{-9}$ | 73.83 |
| $R_g$ (Å) [from GPA] | 42.41 | 43.31 | 29.84 | 31.97 | 29.43 |
| E.R. | 2.60 | 3.18 | 3.43 | 2.63 | 1.84 |
| $D_{max}$ (Å) | ~172 | ~170 | ~115 | ~112 | ~110 |
| Porod volume estimate, $V_p$ ($Å^3$) | 175299 | 191419 | 66194 | 68670 | 101631 |

| | | | | | |
|---|---|---|---|---|---|
| Porod exponent | 3.5 | 3.5 | 3.7 | 3.4 | 3.0 |
| **Molecular mass determination** | | | | | |
| MM (kDa) [from *SAXSMoW* on final merged curve] | 100.80 ($q$ = 0.35 Å$^{-1}$) | 107.10 ($q$ = 0.40 Å$^{-1}$) | 42.90 ($q$ = 0.40 Å$^{-1}$) | 39.20 ($q$ = 0.30 Å$^{-1}$) | 50.60 ($q$ = 0.45 Å$^{-1}$) |
| MM (kDa) [from $Q_R$ on final merged curve] | 96.11 ($q$ = 0.35 Å$^{-1}$) | 101.56 ($q$ = 0.45 Å$^{-1}$) | 40.12 ($q$ = 0.45 Å$^{-1}$) | 40.62 ($q$ = 0.35 Å$^{-1}$) | 50.96 ($q$ = 0.40 Å$^{-1}$) |
| MM (kDa) [from $V_p$/1.7] | 103.17 | 112.60 | 38.94 | 40.39 | 59.78 |
| MM (kDa) [from *Size & Shape*] | 134.45 | 129.18 | 38.24 | 44.12 | 57.72 |
| MM (kDa) [from *Bayesian Inference*] | 94.25 | 109.08 | 38.53 | 34.58 | 50.86 |
| Calculated MM from sequence (kDa) | 98.25 | 99.01 | 38.39 | 40.03 | 49.81 |
| **Ensemble modeling** | | | | | |
| Conformer generation and selection | BILBOMD | BILBOMD | BILBOMD | BILBOMD | BILBOMD |
| Protein regions selected to be fixed/rigid | 22-388, 412-441 | 28-383, 401-449 | 42-95,105-150,164-172,208-222,244-288 | 40-304,307-314,329-349 | 22-257, 272-317, 339-370 |
| **Software employed** | | | | | |
| Data processing and analysis | ATSAS, RAW | ATSAS, RAW | ATSAS, RAW | ATSAS, RAW | ATSAS, RAW |
| Computation of theoretical intensities and fitting | FoXS, MultiFoXS | FoXS, MultiFoXS | FoXS, MultiFoXS | FoXS, MultiFoXS | FoXS, MultiFoXS |
| **SASDBD entry** | SASDTA5 | SASDTB5 | SASDT75 | SASDT85 | SASDT95 |

Abbreviations: *I(0)*, extrapolated scattering intensity at zero angle; $R_g$, radius of gyration calculated using either Guinier approximation (from Guinier) or the indirect Fourier transform package GNOM [from *p(r)*]; *MM*, molecular mass; $D_{max}$, maximal particle dimension; $V_p$, Porod volume; $V_{ex}$, particle excluded volume; E.R., elongation ratio

*Momentum transfer $|q| = 4\pi\sin(\theta)/\lambda$

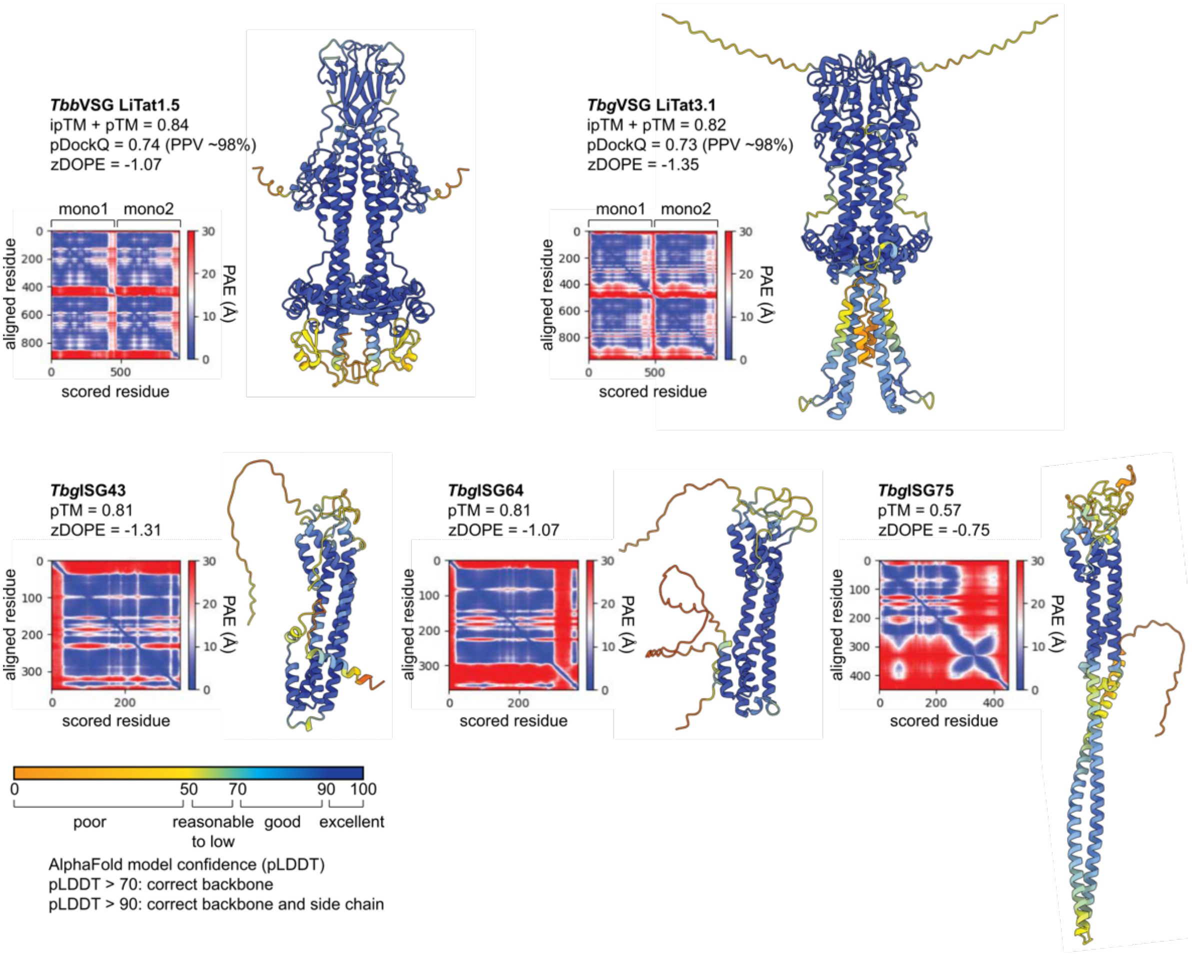

**Supplementary Figure 1. Overview of the AlphaFold2 prediction models and associated quality metrics.** Cartoon representations of the models generated by AlphaFold2 (*Tbg*ISG43, *Tbg*ISG64, and *Tbg*ISG75) and AlphaFold-Multimer (*Tbb*VSG LiTat1.5 and *Tbg*VSGLiTat3.1). The models are colored according to the predicted local distance difference test (pLDDT) score, which reflects (local) model quality as indicated by the legend at the bottom. For all structures, the predicted aligned error (PAE), the normalized discrete optimized protein energy (zDOPE), and overall predicted template modeling (pTM) scores are also shown. The PAE provides a distance error for every residue pair, and is calculated for each residue x (scored residue) when the predicted and true structures are aligned on residue y (aligned residue). A zDOPE < -1 indicates that the distribution of atom pair distances in the model resembles that found in a large sample of known protein structures and that at least 80% of the model's Cα atoms are within 3.5 Å of their correct positions. The pTM (score between 0 and 1) provides a measure of similarity between two protein structures (in this case, the predicted and unknown true structure) over all residues and thus reports on the accuracy of prediction within a single chain. For multimers, the pDockQ and AlphaFold-Multimer model confidence (0.8*ipTM + 0.2*pTM) are also shown. The interface pTM (ipTM, score between 0 and 1) provides a measure of similarity between two protein structures (in this case, the predicted and unknown true structure) over only interfacing residues and thus reports on the accuracy of prediction for a complex. The pDockQ score (between 0 and 1) is another confidence metric for protein complexes that takes into account the number of interfacing residues and their pLDDT scores. Determination of the pDockQ score can be associated to a positive predictive value (PPV), which provides an estimate for the probability that the solution is a true positive.

#### *Tbg*ISG43

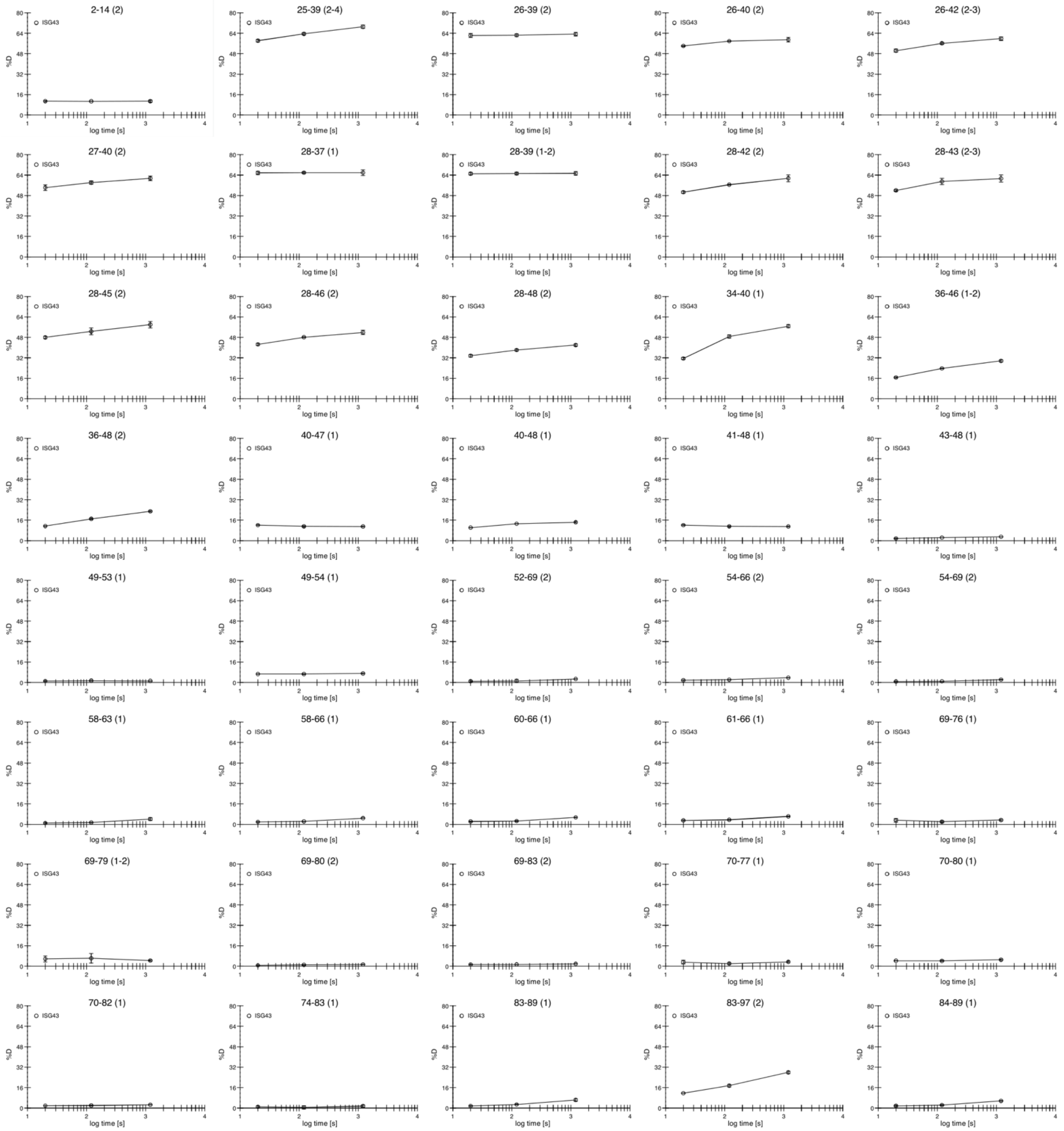
2-14 (2)
25-39 (2-4)
26-39 (2)
26-40 (2)
26-42 (2-3)
27-40 (2)
28-37 (1)
28-39 (1-2)
28-42 (2)
28-43 (2-3)
28-45 (2)
28-46 (2)
28-48 (2)
34-40 (1)
36-46 (1-2)
36-48 (2)
40-47 (1)
40-48 (1)
41-48 (1)
43-48 (1)
49-53 (1)
49-54 (1)
52-69 (2)
54-66 (2)
54-69 (2)
58-63 (1)
58-66 (1)
60-66 (1)
61-66 (1)
69-76 (1)
69-79 (1-2)
69-80 (2)
69-83 (2)
70-77 (1)
70-80 (1)
70-82 (1)
74-83 (1)
83-89 (1)
83-97 (2)
84-89 (1)
ISG43
%D
log time [s]

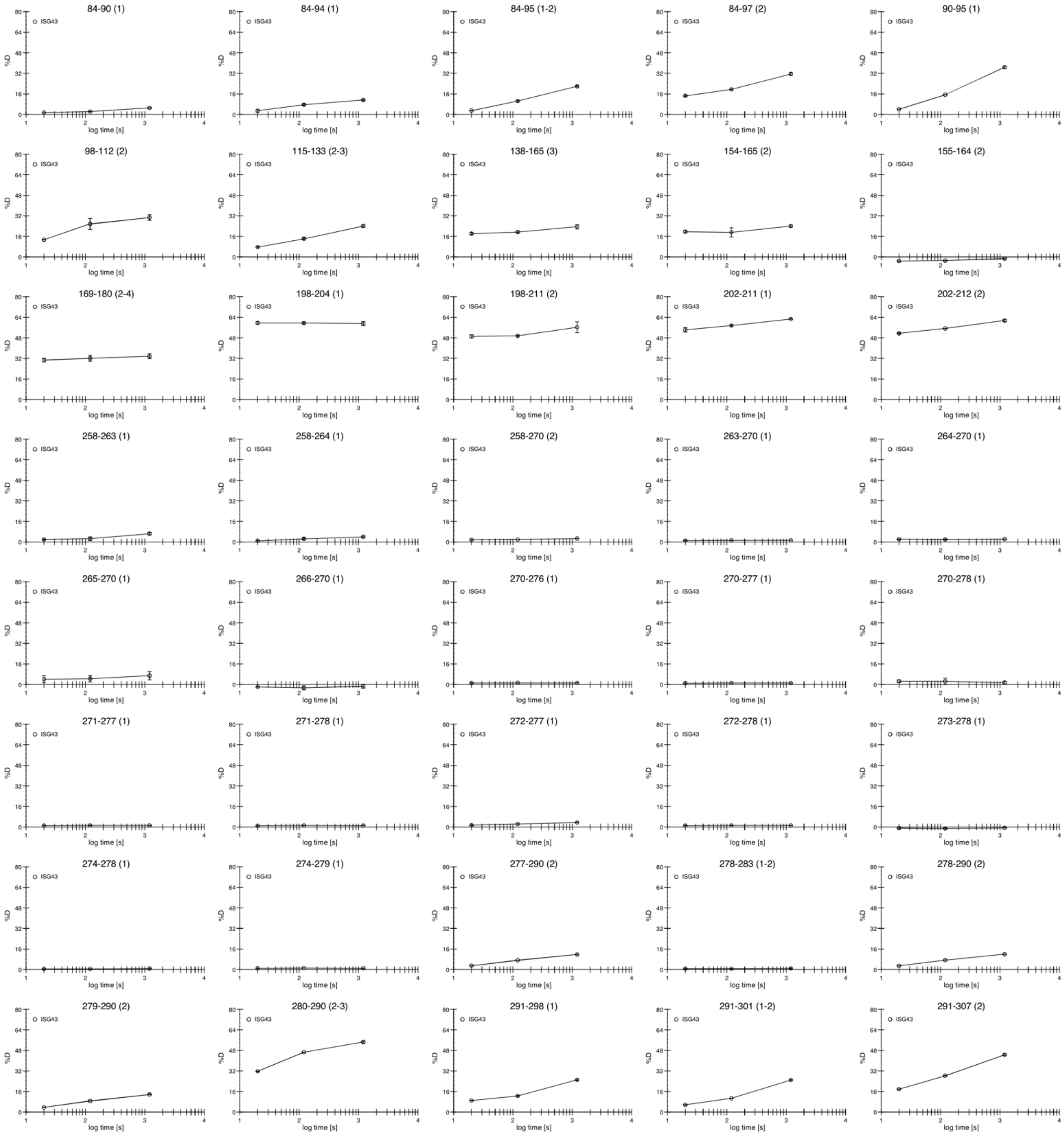
84-90 (1)
84-94 (1)
84-95 (1-2)
84-97 (2)
90-95 (1)
98-112 (2)
115-133 (2-3)
138-165 (3)
154-165 (2)
155-164 (2)
169-180 (2-4)
198-204 (1)
198-211 (2)
202-211 (1)
202-212 (2)
258-263 (1)
258-264 (1)
258-270 (2)
263-270 (1)
264-270 (1)
265-270 (1)
266-270 (1)
270-276 (1)
270-277 (1)
270-278 (1)
271-277 (1)
271-278 (1)
272-277 (1)
272-278 (1)
273-278 (1)
274-278 (1)
274-279 (1)
277-290 (2)
278-283 (1-2)
278-290 (2)
279-290 (2)
280-290 (2-3)
291-298 (1)
291-301 (1-2)
291-307 (2)
ISG43
%D
log time [s]

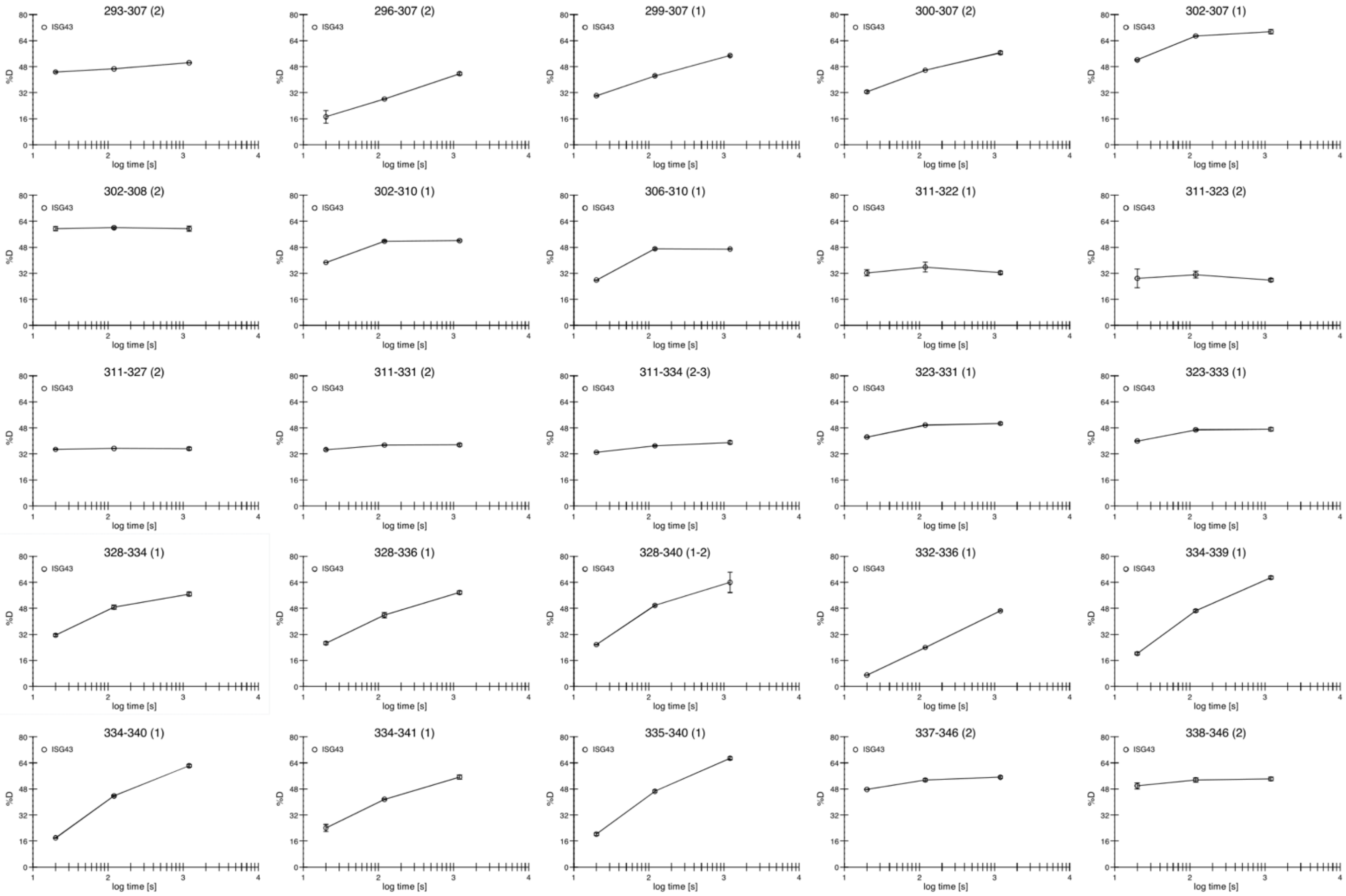
293-307 (2)
296-307 (2)
299-307 (1)
300-307 (2)
302-307 (1)
302-308 (2)
302-310 (1)
306-310 (1)
311-322 (1)
311-323 (2)
311-327 (2)
311-331 (2)
311-334 (2-3)
323-331 (1)
323-333 (1)
328-334 (1)
328-336 (1)
328-340 (1-2)
332-336 (1)
334-339 (1)
334-340 (1)
334-341 (1)
335-340 (1)
337-346 (2)
338-346 (2)
ISG43
%D
log time [s]
0
16
32
48
64
80
1
2
3
4

### *Tbg*ISG64

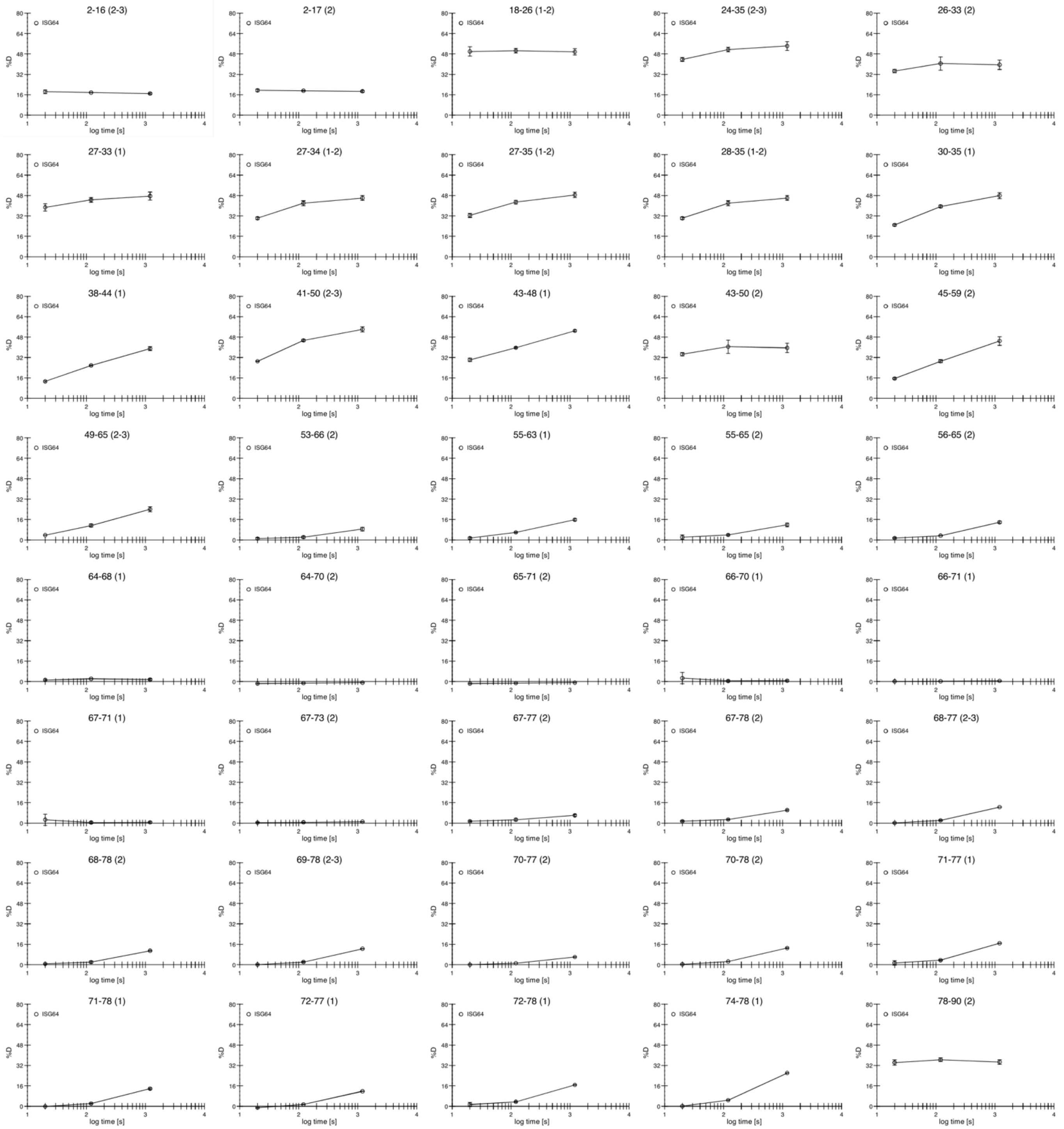
2-16 (2-3)
2-17 (2)
18-26 (1-2)
24-35 (2-3)
26-33 (2)
27-33 (1)
27-34 (1-2)
27-35 (1-2)
28-35 (1-2)
30-35 (1)
38-44 (1)
41-50 (2-3)
43-48 (1)
43-50 (2)
45-59 (2)
49-65 (2-3)
53-66 (2)
55-63 (1)
55-65 (2)
56-65 (2)
64-68 (1)
64-70 (2)
65-71 (2)
66-70 (1)
66-71 (1)
67-71 (1)
67-73 (2)
67-77 (2)
67-78 (2)
68-77 (2-3)
68-78 (2)
69-78 (2-3)
70-77 (2)
70-78 (2)
71-77 (1)
71-78 (1)
72-77 (1)
72-78 (1)
74-78 (1)
78-90 (2)
ISG64
%D
log time [s]

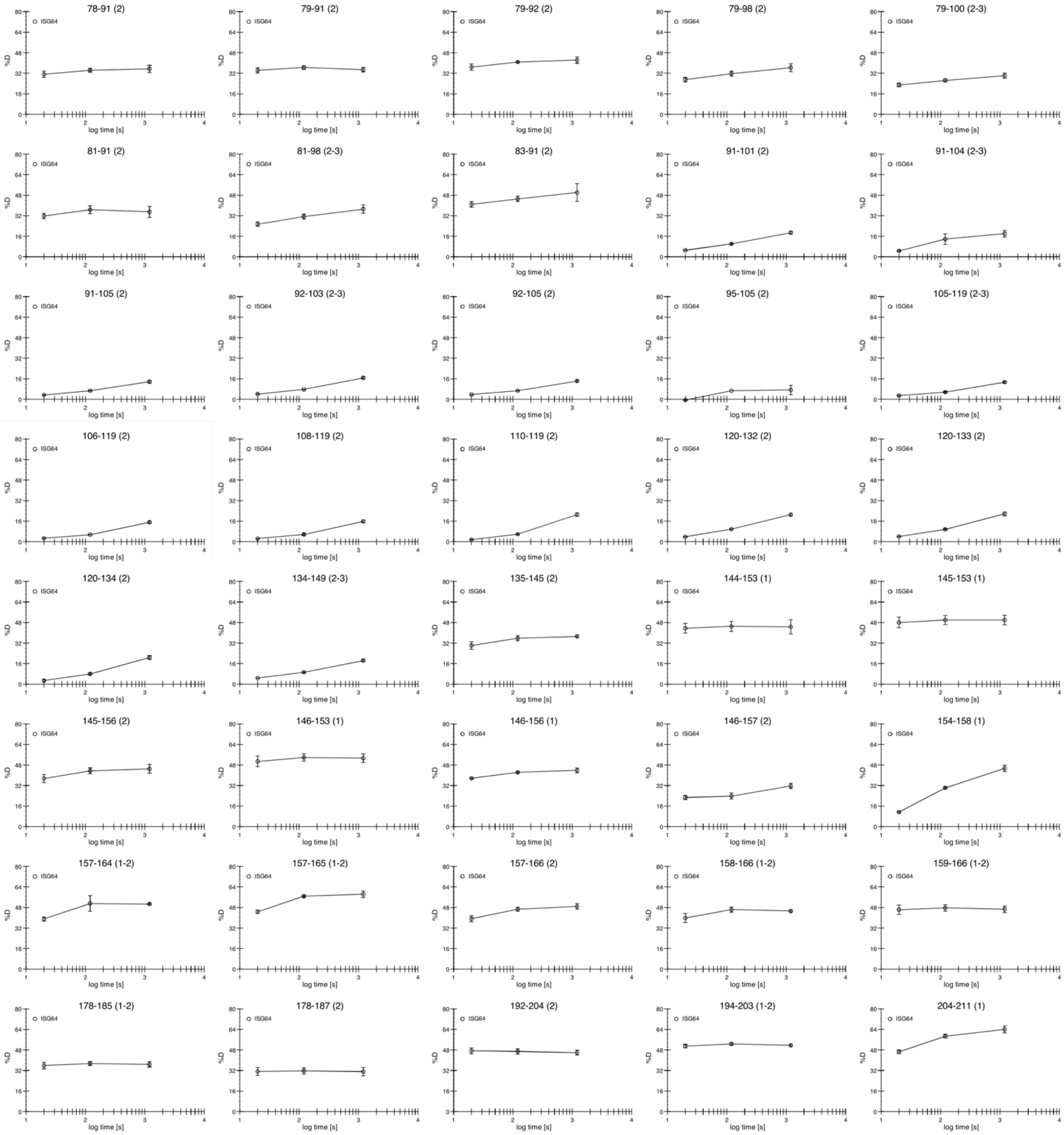
78-91 (2)
79-91 (2)
79-92 (2)
79-98 (2)
79-100 (2-3)
81-91 (2)
81-98 (2-3)
83-91 (2)
91-101 (2)
91-104 (2-3)
91-105 (2)
92-103 (2-3)
92-105 (2)
95-105 (2)
105-119 (2-3)
106-119 (2)
108-119 (2)
110-119 (2)
120-132 (2)
120-133 (2)
120-134 (2)
134-149 (2-3)
135-145 (2)
144-153 (1)
145-153 (1)
145-156 (2)
146-153 (1)
146-156 (1)
146-157 (2)
154-158 (1)
157-164 (1-2)
157-165 (1-2)
157-166 (2)
158-166 (1-2)
159-166 (1-2)
178-185 (1-2)
178-187 (2)
192-204 (2)
194-203 (1-2)
204-211 (1)
ISG64
%D
log time [s]

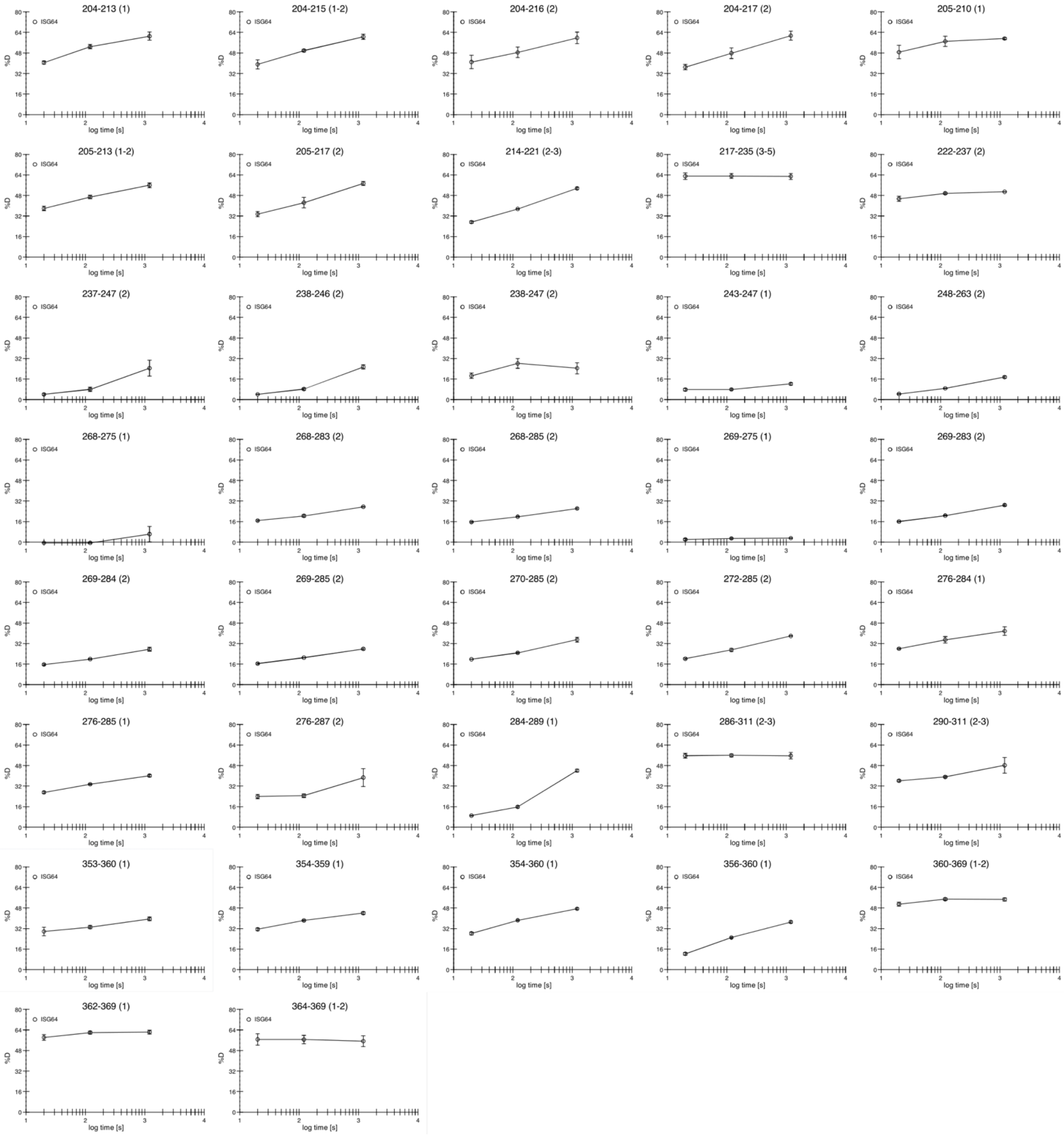
204-213 (1)
204-215 (1-2)
204-216 (2)
204-217 (2)
205-210 (1)
205-213 (1-2)
205-217 (2)
214-221 (2-3)
217-235 (3-5)
222-237 (2)
237-247 (2)
238-246 (2)
238-247 (2)
243-247 (1)
248-263 (2)
268-275 (1)
268-283 (2)
268-285 (2)
269-275 (1)
269-283 (2)
269-284 (2)
269-285 (2)
270-285 (2)
272-285 (2)
276-284 (1)
276-285 (1)
276-287 (2)
284-289 (1)
286-311 (2-3)
290-311 (2-3)
353-360 (1)
354-359 (1)
354-360 (1)
356-360 (1)
360-369 (1-2)
362-369 (1)
364-369 (1-2)
ISG64
%D
log time [s]

#### ***Tbg*ISG75**

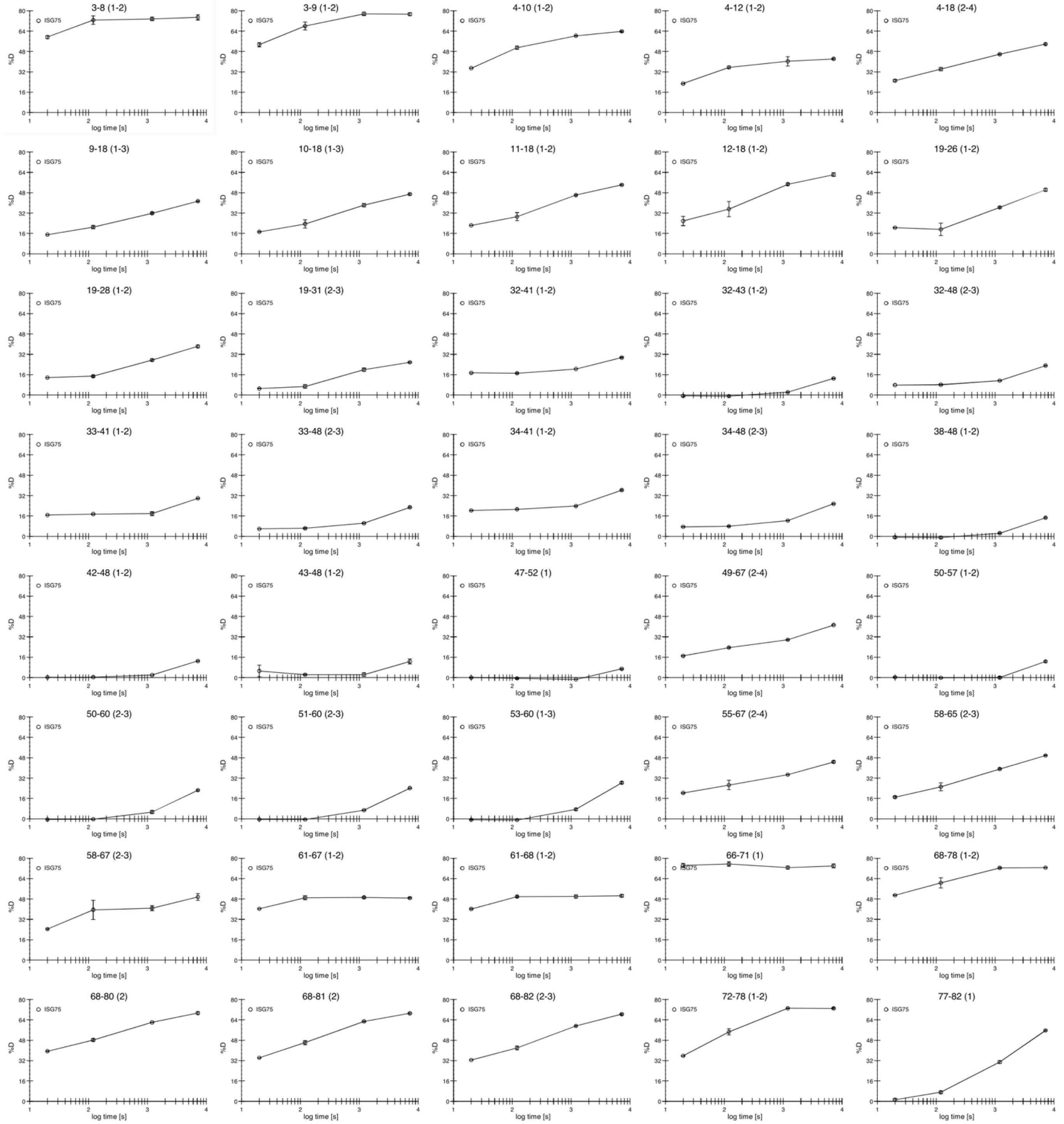
3-8 (1-2)
3-9 (1-2)
4-10 (1-2)
4-12 (1-2)
4-18 (2-4)
9-18 (1-3)
10-18 (1-3)
11-18 (1-2)
12-18 (1-2)
19-26 (1-2)
19-28 (1-2)
19-31 (2-3)
32-41 (1-2)
32-43 (1-2)
32-48 (2-3)
33-41 (1-2)
33-48 (2-3)
34-41 (1-2)
34-48 (2-3)
38-48 (1-2)
42-48 (1-2)
43-48 (1-2)
47-52 (1)
49-67 (2-4)
50-57 (1-2)
50-60 (2-3)
51-60 (2-3)
53-60 (1-3)
55-67 (2-4)
58-65 (2-3)
58-67 (2-3)
61-67 (1-2)
61-68 (1-2)
66-71 (1)
68-78 (1-2)
68-80 (2)
68-81 (2)
68-82 (2-3)
72-78 (1-2)
77-82 (1)
ISG75
%D
log time [s]

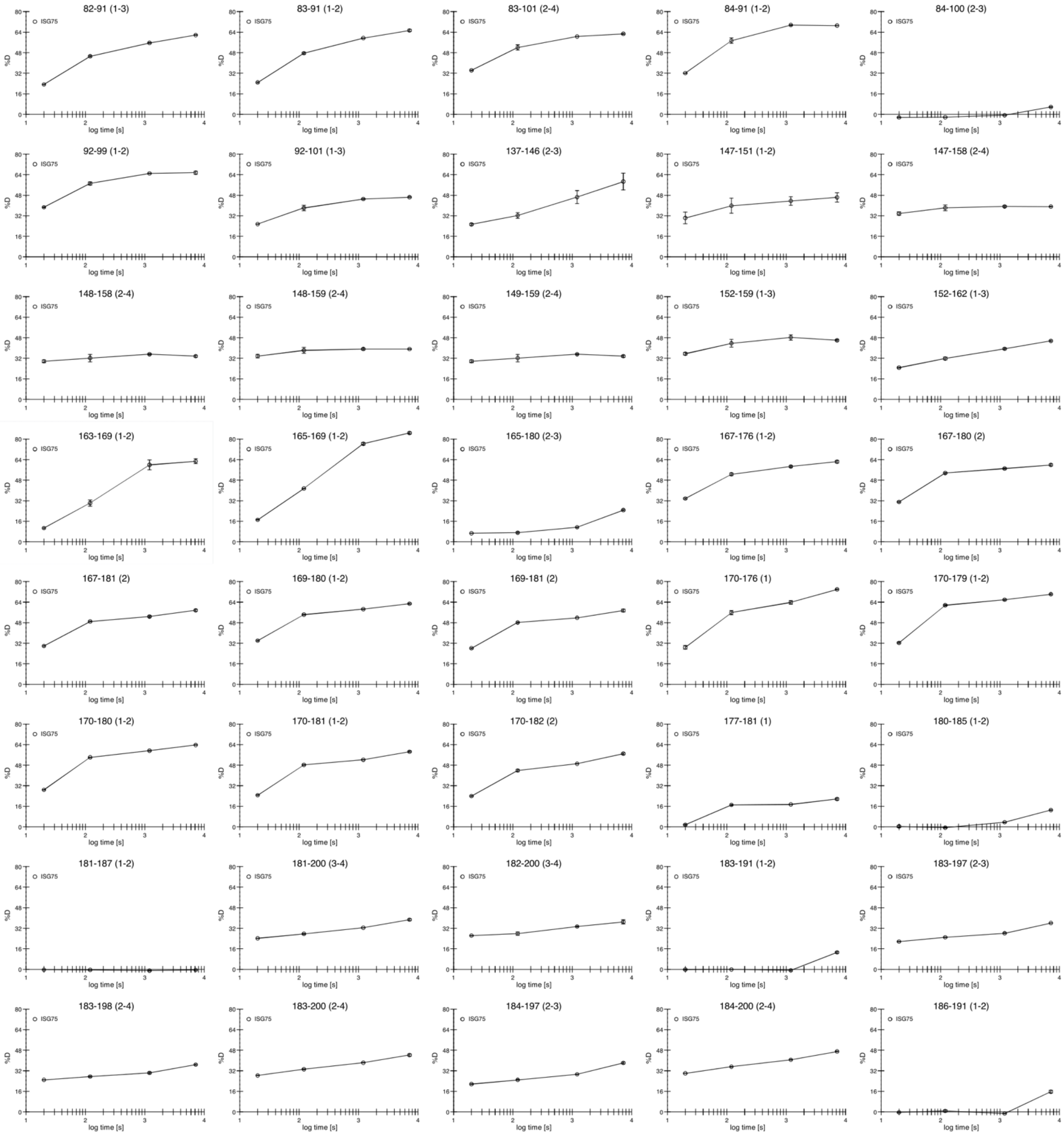
82-91 (1-3)
83-91 (1-2)
83-101 (2-4)
84-91 (1-2)
84-100 (2-3)
92-99 (1-2)
92-101 (1-3)
137-146 (2-3)
147-151 (1-2)
147-158 (2-4)
148-158 (2-4)
148-159 (2-4)
149-159 (2-4)
152-159 (1-3)
152-162 (1-3)
163-169 (1-2)
165-169 (1-2)
165-180 (2-3)
167-176 (1-2)
167-180 (2)
167-181 (2)
169-180 (1-2)
169-181 (2)
170-176 (1)
170-179 (1-2)
170-180 (1-2)
170-181 (1-2)
170-182 (2)
177-181 (1)
180-185 (1-2)
181-187 (1-2)
181-200 (3-4)
182-200 (3-4)
183-191 (1-2)
183-197 (2-3)
183-198 (2-4)
183-200 (2-4)
184-197 (2-3)
184-200 (2-4)
186-191 (1-2)
ISG75
%D
log time [s]

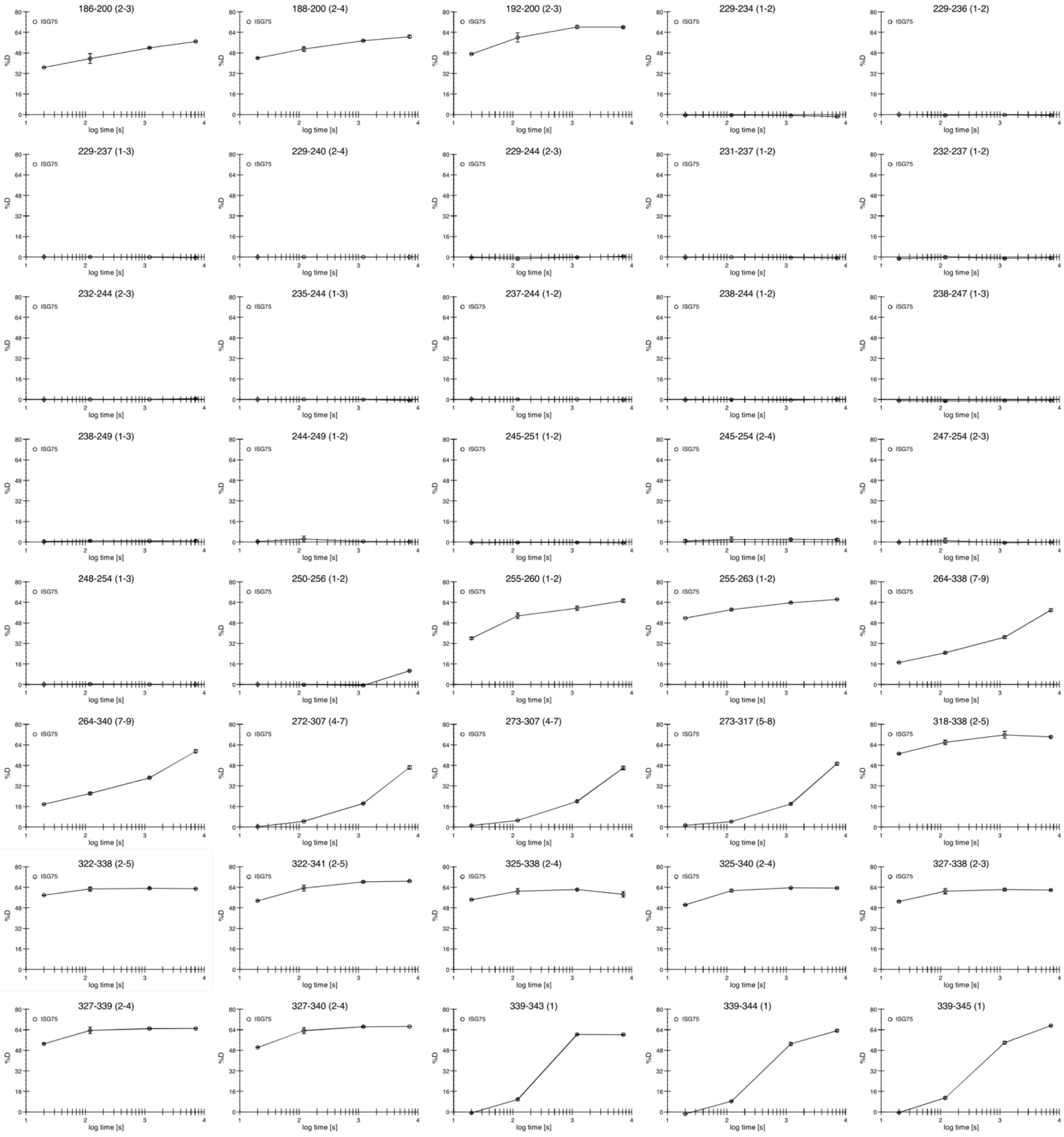
186-200 (2-3)
188-200 (2-4)
192-200 (2-3)
229-234 (1-2)
229-236 (1-2)
229-237 (1-3)
229-240 (2-4)
229-244 (2-3)
231-237 (1-2)
232-237 (1-2)
232-244 (2-3)
235-244 (1-3)
237-244 (1-2)
238-244 (1-2)
238-247 (1-3)
238-249 (1-3)
244-249 (1-2)
245-251 (1-2)
245-254 (2-4)
247-254 (2-3)
248-254 (1-3)
250-256 (1-2)
255-260 (1-2)
255-263 (1-2)
264-338 (7-9)
264-340 (7-9)
272-307 (4-7)
273-307 (4-7)
273-317 (5-8)
318-338 (2-5)
322-338 (2-5)
322-341 (2-5)
325-338 (2-4)
325-340 (2-4)
327-338 (2-3)
327-339 (2-4)
327-340 (2-4)
339-343 (1)
339-344 (1)
339-345 (1)
ISG75
%D
log time [s]

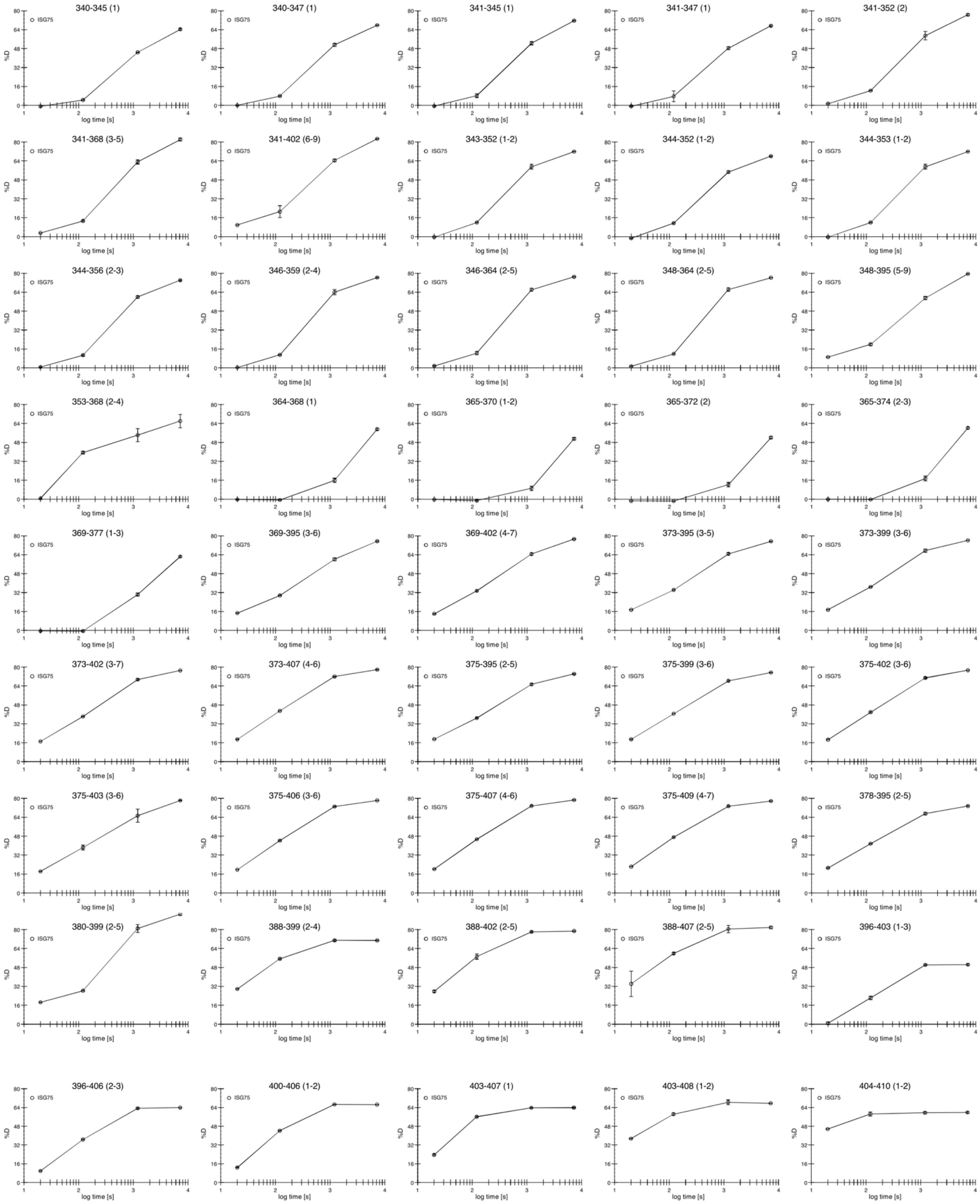

**Supplementary Figure 2. HDX-MS analysis of *Tbg*ISGs.** Hydrogen/deuterium exchange plots for *T. b. gambiense* ISG43, ISG64 and ISG75. All time points were measured in triplicates. Figures were generated with MSTools [1].
